# Structure-forming CAG/CTG repeats interfere with gap repair to cause repeat expansions and chromosome breaks

**DOI:** 10.1101/2022.03.22.485386

**Authors:** Erica J. Polleys, Isabella Del Priore, Marjorie de la Rosa Mejia, James E. Haber, Catherine H. Freudenreich

## Abstract

Expanded CAG/CTG repeats are sites of DNA damage, leading to changes in repeat length. To determine how ssDNA gap filling affects repeat instability, we inserted (CAG)_70_ or (CTG)_70_ repeats into a single-strand annealing (SSA) assay system such that resection and filling in the ssDNA gap would occur across the repeat tract. After resection, when the CTG sequence was the single-stranded template for fill-in synthesis, repeat contractions were elevated and the ssDNA created a fragile site that led to large deletions involving flanking homologous sequences. In contrast, resection was inhibited when CTG was on the resected strand, resulting in repeat expansions. Deleting Rad9, the ortholog of 53BP1, rescued repeat instability and lost viability by increasing resection and fill-in speed. Deletion of Rad51 increased CTG contractions and decreased survival, implicating Rad51 in protecting ssDNA during gap filling. Taken together, DNA sequence within a single-stranded gap determines repair kinetics, fragility, and repeat instability.

## Introduction

Alternative DNA structures formed by expanded CAG/CTG repeats can result in the formation of a barrier, making it difficult for both DNA replication and repair machineries to proceed smoothly. As such, these repetitive regions are hotspots for genomic change. When a breakage event occurs within a CAG/CTG repeat tract, it can be repaired by homologous recombination (HR), using a region of homology on a sister chromatid or homologous chromosome as a template. Though HR is generally considered to be an error-free mechanism of repair, the fidelity of repair through a CAG/CTG repeat may be compromised, as HR has been shown to be a mechanism for repeat expansions^1^. Additionally, recombination-dependent expansions and contractions are seen in strains carrying mutations in proteins important in DNA repair or replication, indicating that CAG repeat length changes occur through HR-dependent mechanisms in cells that are experiencing replication stress or DNA damage^2^.

The mechanisms that drive repeat instability during HR are not fully understood. Key steps in HR involve the 5’ to 3’ resection of DSB ends and the subsequent filling in of ssDNA regions. Filling in ssDNA gaps is one point where CAG/CTG repeats could expand, as polymerases may be prone to slippage across the CAG/CTG repeat. Repair DNA synthesis is 1000-fold more mutagenic than replication of the same sequences^3, 4^. Polymerase slippage while filling in ssDNA gaps that arise during mismatch repair (MMR) was proposed to be responsible for germline repeat expansions in a Huntington’s disease mouse model^5^, but how gap filling proceeds in the context of a CAG/CTG repeat has not been directly determined. The processivity of the polymerase, size of the gap and stability of the DNA secondary structure could all contribute to trinucleotide repeat (TNR) instability during gap repair^6^.

Resection is a highly conserved process that is considered one of the key steps that drives repair away from end-joining and toward HR. Recently, it has been shown that G4 quadruplex sequences are a barrier to resection in mammalian cells, and require the helicase PIF1, along with BRCA1, to resolve the secondary structure and promote recombinational repair^7^. Resection is restrained by 53BP1^8–10^ and it has been proposed that the 53BP1 homolog in budding yeast, Rad9, forms a dynamic barrier at the ssDNA/dsDNA junction through interaction with Dpb11 and the phosphorylated form of histone H2A, γH2AX^11^. In a *rad9*Δ mutant there is increased ssDNA and enrichment of the single-strand binding protein complex, RPA, as well as the Rad51 recombinase and the mediator of Rad51 assembly, Rad52, around the site of a targeted double-strand break (DSB)^12^. Outside of its role in resection, Rad9 has a well-characterized role in activation of the DNA damage checkpoint^13^. Previous work has shown that loss of Rad9 resulted in a significant increase in CAG repeat fragility and instability due to its role in checkpoint activation^14^. It remained to be determined whether repeat stability would be impacted by resection kinetics, and if so, whether Rad9 played a role.

To determine whether ssDNA gap repair results in CAG/CTG repeat instability, we used an assay system that repairs an induced DSB via single strand annealing (SSA)^15^ and modified it such that ssDNA gap filling would occur through a long CAG/CTG repeat tract. We found that the template of the repeat dictates the efficiency of repair kinetics as well as the type and magnitude of repeat instability. When a (CAG)_70_ repeat tract was the template for gap filling we noted no apparent loss in expected repair, though resection was impaired and small-scale repeat expansions were observed. Conversely, when a (CTG)_70_ repeat tract was the ssDNA template for filling in, there was a significant decrease in canonical repair, caused by breakage at the repeat tract; the recovered repair products contained large scale deletions and repeat contractions. Increasing the rate of resection and thus the kinetics of RPA and Rad51 loading by deleting Rad9 reduced repeat contractions and increased viability. In contrast, deleting Rad51 resulted in increased repeat contractions, suggesting that Rad51 can function like RPA in preventing DNA secondary structure formation at expanded repeat tracts. For the (CTG)_70_ template, loss of Rad51 also resulted in persistent ssDNA near the repeat tract due to loss of break-induced replication (BIR) as a backup repair pathway leading to further decreases in survival. We propose that resection and gap filling through a repeat tract are key steps required to prevent repeat instability and protect genome integrity. This work illustrates that large ssDNA gaps create an ideal environment for DNA secondary structure formation, which can act as a fragile site to cause large scale deletions. Therefore, the repeat content and structure-forming potential of the region surrounding a DSB determines repair pathway choice and repair fidelity.

## Results

### Identity of the repeat on the template strand determines survival during gap fill-in

To determine whether fill-in synthesis resulted in CAG/CTG repeat instability, we integrated a (CAG/CTG)_70_ repeat tract into a strain that repairs an induced DSB via SSA^15^. A single HO endonuclease induced DSB in the *LEU2* gene results in resection on both sides of the DSB. Once 25 kb of resection occurs, the more distant U2 homologous region is exposed and repair via SSA occurs. Finally, regions that were rendered single-stranded during resection are filled in by DNA polymerases (Figure 1a). Successful repair via SSA is measured as percent viability. To study instability of the CAG/CTG repeat during fill-in synthesis, TNR tracts were integrated ∼13 kb away from the DSB on the centromere-proximal side. As resection occurs on both sides of the break^16^, the resected DNA containing the repeat tract will be a site for polymerase-mediated filling in after annealing of the U2 regions and clipping of the non- homologous tails^17^. We inserted repeat tracts such that either 70 CAG or CTG repeats were on the strand that is not resected and thus serves as the template for DNA fill-in synthesis (Figure 1b).

**Figure 1.**
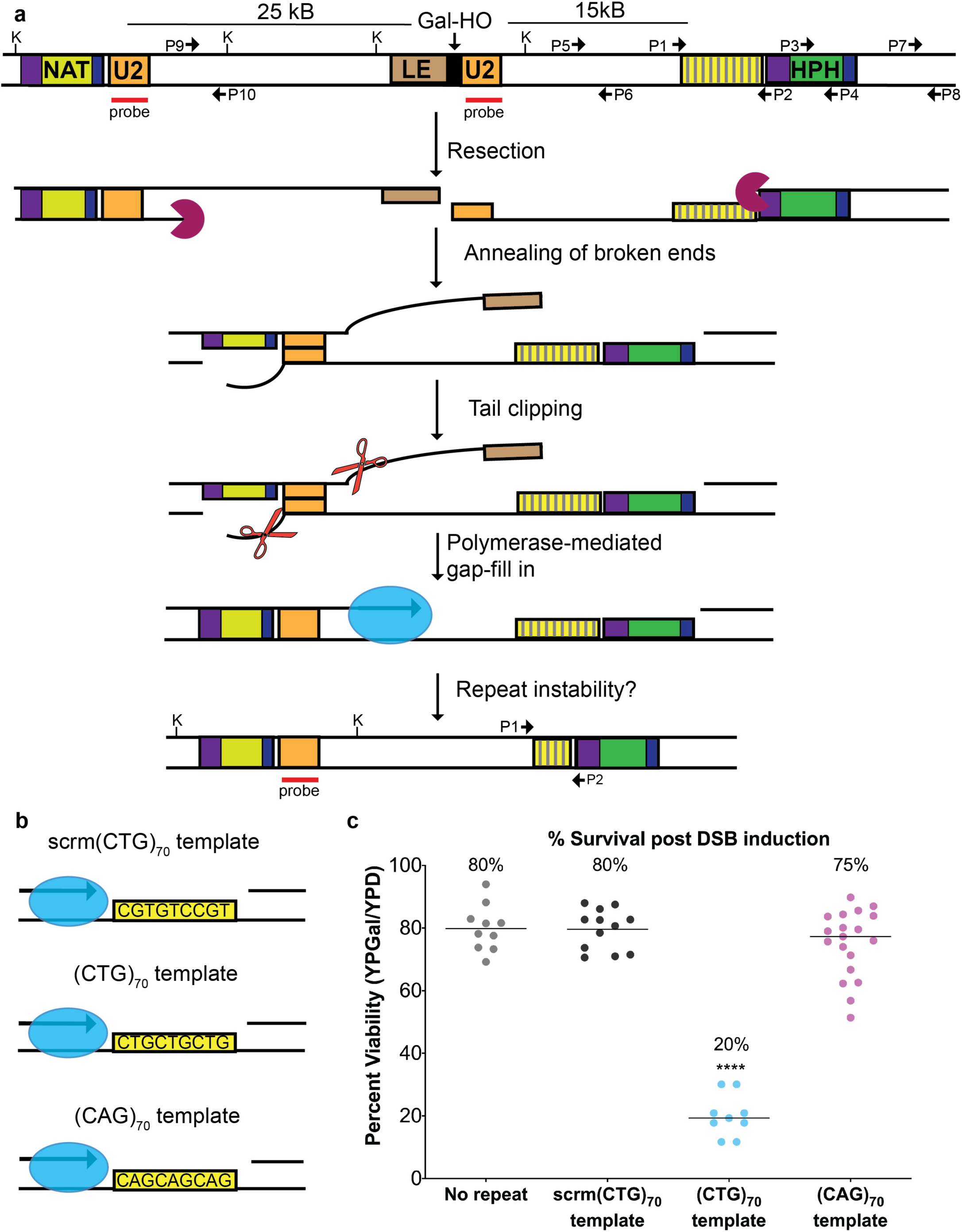
Template of the repeat tract determines survival post-DSB induction. **a)** Assay system to study gap repair mediated CAG/CTG repeat instability. Modifying strain YMV80 ^15^, we inserted a CAG/CTG repeat tract ∼13 kb centromere-proximal from an HO endonuclease cut site within *LEU2*. Following galactose-induction of the HO endonuclease, repair via SSA occurs between the two 1-kb U2 homologies. Fill-in synthesis occurs through the inserted repeat tract after U2 annealing. Relevant *Kpn*I sites are marked. Probe used for analysis of SSA repair is marked. P1-P10 primer locations shown; sequences in Table 1. **b)** In this assay system, the template of the CAG/CTG repeat is defined as the sequence that remains after 5’ to 3’ resection of the DNA creates a single-stranded template. The fill-in template of the inserted repeat tract is either a scrambled control (scrm(CTG)_70_), (CTG)_70_ or (CAG)_70_. **c)** Percent viability of the no repeat control (n=10), scrambled (n=12), (CTG)_70_ (n=9), and (CAG)_70_ (n=19) template strains. Statistical significance between (CTG)_70_ compared to the scrambled control using the Student’s t-test (p<0.0001).

To be able to attribute any altered outcomes to the presence of the repeat tract, we created a scrambled control containing a non-structure-forming sequence with equal amounts of C, T, and G on the template (scrm(CTG)_70_) (Figure 1b). The scrm(CTG)_70_ control has similar viability compared to the original assay strain (no repeat) (Figure 1c)^15^. As discussed in more detail below, monitoring of HO cleavage, 5’ to 3’ resection and product formation, as well as the activation and persistence of the DNA damage checkpoint (i.e. phosphorylation of Rad53) showed no differences between the no repeat and scrm(CTG)_70_ strains (SFigure1 a, b & c); thus the addition of a non-repetitive sequence does not alter repair and is a satisfactory control for our repeat-containing strains. Interestingly, strains that had an inserted (CTG)_70_ repeat on the fill-in template showed a significant four-fold decrease in viability compared to the scrambled control strain (Figure 1c). In contrast, strains that had a (CAG)_70_ fill-in template had no viability defect. The differences in viability between the (CTG)_70_ and (CAG)_70_ strains suggested that the template of an expanded repeat tract could influence repair outcome, and thus cell survival.

### Identity of the sequence on the template strand determines resection kinetics and repair efficiency of gap filling

We next explored whether the addition of an expanded repeat tract altered the kinetics or efficiency of resection and repair, and if this could explain the loss in viability seen in the (CTG)_70_ template strain. Tracking DSB induction and repair product formation via Southern blotting^15^ revealed that the expected SSA repair product still forms in the (CTG)_70_ template strain with the same timing, though at a reduced level, compared to the scrambled control or the (CAG)_70_ template strain (Figure 2a). One possible reason for the decreased viability could be due to persistent activation of the DNA damage checkpoint^15^; therefore, we assessed the state of Rad53 phosphorylation by Western blot after DSB induction. None of the strains showed persistent hyperactivation of Rad53, ruling out that the decreased viability in the (CTG)_70_ template strain was due to a defect in recovery from checkpoint activation (SFigure 1c).

**Figure 2.**
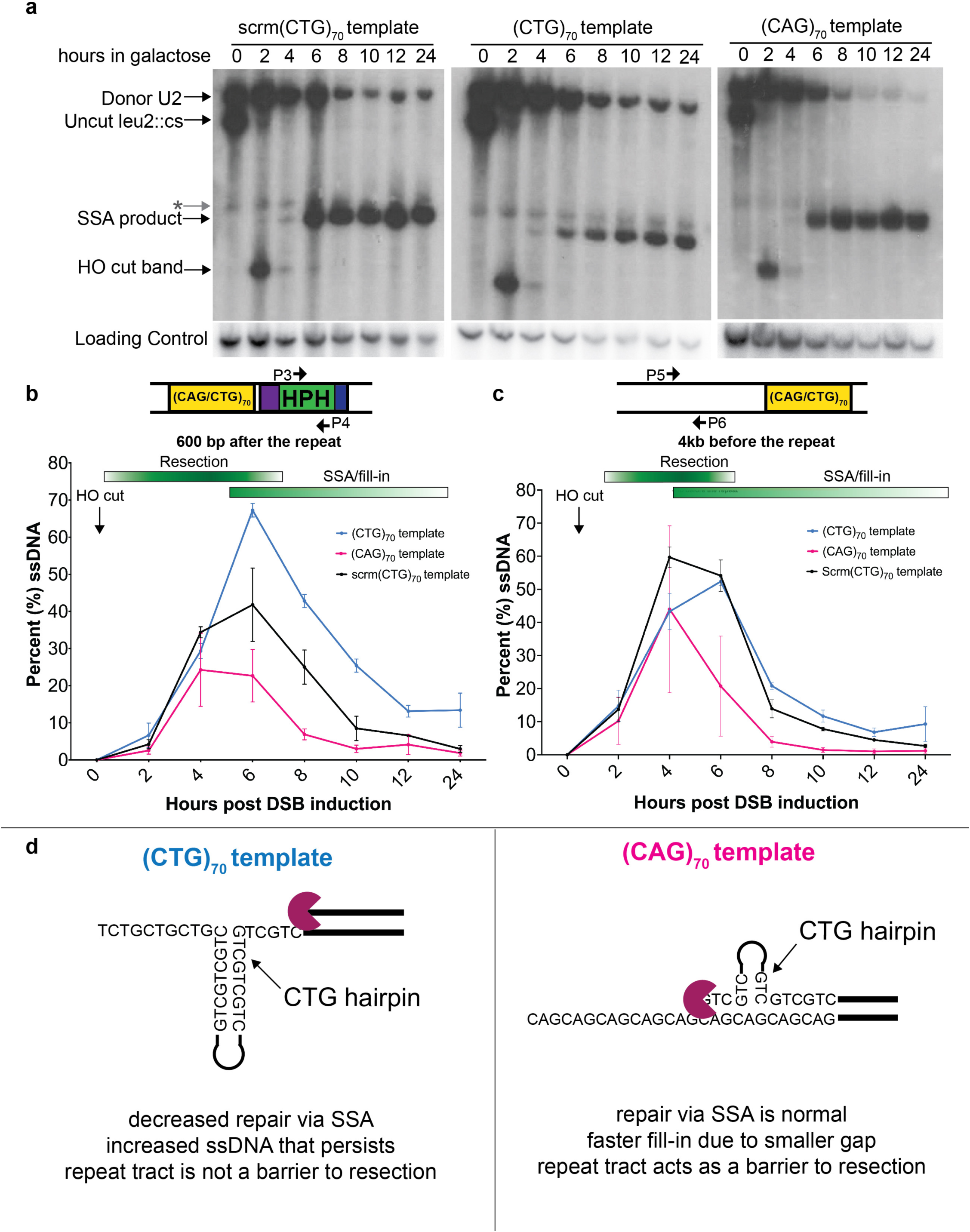
Template of the repeat results in alterations in repair kinetics during gap filling. **a)** Southern blot analysis after addition of 2% galactose to induce a DSB within *LEU2*. Representative Southern blot is shown; number of replicates: scrm(CTG)_70_ n=3, (CTG)_70_ n=4, (CAG)_70_ n=5. Probe location is marked in Figure 1a; loading control probe is to a portion of the *TRA1* gene. Non-specific hybridization of the probe is marked with a gray star. **b)** Formation and disappearance of ssDNA proximal to the repeat tract, as described in Methods, using primers P3 and P4. Number of replicates: scrm(CTG)_70_ n=3, (CTG)_70_ n=4 and (CAG)_70_ n=3. **c)** ssDNA abundance in a region 4 kb before the repeat locus as described above, with primers P5 and P6. Number of replicates: scrm(CTG)_70_ n=3, (CTG)_70_ n=3 and (CAG)_70_ n=2. **d)** Summary of results and interpretation for resection and secondary structure formation in each template.

As this assay relies on extensive resection to expose the homologous repair sequence (U2, Figure 1a), it is an ideal system to monitor resection and subsequent gap filling. Using a qPCR-based assay to quantify levels of ssDNA^18, 19^, we monitored resection and fill-in kinetics 600 bp after the repeat tract (Figure 2b, primers P3 & P4). In the scrambled control strain, we saw increasing amounts of ssDNA appear between hours 2-6 and a subsequent decrease in ssDNA signal from hours 8-24 (Figure 2b, black line). Though resection across this region is maximal between 2-4 hours, double stranded repair product formation can already be visualized 4-6 hours post-DSB induction (Figure 2a). Therefore, resection, annealing, and fill-in synthesis are likely occurring concurrently. Notably, there is an increase in the amount of ssDNA in the (CTG)_70_ template strain compared to the scrambled control 6 hours post DSB induction; this increased ssDNA remained elevated at all remaining time points (Figure 2b, compare blue and black lines).

During resection, RPA is recruited to ssDNA and helps prevent DNA secondary structure formation^20^. Though this assay system predominantly repairs via SSA, which is Rad51 independent, repair can also occur via BIR which is Rad51-dependent^21^. We monitored enrichment of RPA and Rad51 in both the (CTG)_70_ and scrm(CTG)_70_ strains with the expectation that RPA and Rad51 enrichment would increase during resection and decrease during gap filling. Indeed, in both strains, maximum enrichment of RPA and Rad51 600 bp after the repeat occurs 6 hours after DSB induction when ssDNA is maximal. Importantly, comparing the (CTG)_70_ tract to the scrm(CTG)_70_ control showed no significant differences in the level of RPA and Rad51 enrichment (SFigure 2 a, b). Taken together, these data suggest that there is a defect in completing gap filling and repair when (CTG)_70_ is the fill-in template that is not due to impaired recruitment of RPA and Rad51.

Even though there is no repair defect in the (CAG)_70_ template strain (Figure 2a), less ssDNA accumulates beyond the repeat tract compared to the scrambled control (Figure 2b, compare pink and black lines). When we examined resection and filling in at a location before the repeat tract, we observed no differences in ssDNA levels in the (CAG)_70_ template strain compared to scrm(CTG)_70_ or (CTG)_70_ strains at 4 hours (Figure 2c, primers P5 & P6). The decreased ssDNA accumulation after passing through the TNR suggests that there may be a repeat-specific barrier to resection when (CAG)_70_ is the fill-in template.

Monitoring repair kinetics and resection data suggested a model for how resection through the repeat tract influences repair and thus survival. In strains that have a (CTG)_70_ fill-in template, there is increased, persistent ssDNA at the repeat tract (Figure 2b) which coincides with a decrease in the expected SSA repair product and survival (Figure 1c, 2a). These results suggest that CTG hairpin formation on the ssDNA template strand is impeding repair (Figure 2d, left panel). In strains that have a (CAG)_70_ fill-in template, we hypothesize that a CTG hairpin is forming on the 5’ recessed end, which is sufficient to disrupt resection (Figure 2d, right panel). Therefore, in the (CAG)_70_ template strain, filling in occurs faster because there was a smaller ssDNA gap due to the hairpin-impaired resection. Taken together, we conclude that a CTG hairpin on the 5’ recessed strand acts as a barrier to resection, decreasing the amount of ssDNA, whereas a CTG hairpin on the template strand impedes or slows gap filling, increasing the amount of ssDNA.

### CAG/CTG expansions and contractions occur during gap filling

Our assay system tracks changes in repeat length both in conditions where there is no induced HO break (in glucose) or post induction of the DSB at *LEU2* (in galactose) as distinct populations. Instability of expanded repeat tracts can occur during normal DNA transactions such as DNA replication and repair. The direction of replication influences the basal stability of structure-forming repeats: CTG is on the lagging strand template favors contractions, and CTG on the nascent lagging strand favors expansions^22–24^ because CTG repeats can form a more thermodynamically stable hairpin compared to CAG repeats^25^. In addition, expanded repeats are fragile and prone to DSBs within the repeat which can result in out of register annealing events that lead to expansions or contractions^2^. Indeed, DSB induction using Cas9 to target an expanded repeat tract increases expansion and contraction frequencies^26, 27^. The observed instability in the no-break condition could be due to either or both of these events. In contrast, the induced HO break condition is testing instability due to DNA synthesis during gap filling which occurs independently of instability due to replication or naturally occurring breaks within the double-stranded repeat.

To determine whether gap filling resulted in changes in TNR repeat stability, we used PCR to compare tract length changes in colonies from the no-break and HO-induced break (DSB) conditions using primers that span the repeat tract (Figure 1a, Primers P1 & P2). Products were separated using capillary gel electrophoresis (Figure 3a, 3b) and the frequency of each PCR product size for independent colonies tested was plotted (Figure 3a, 3b).

**Figure 3.**
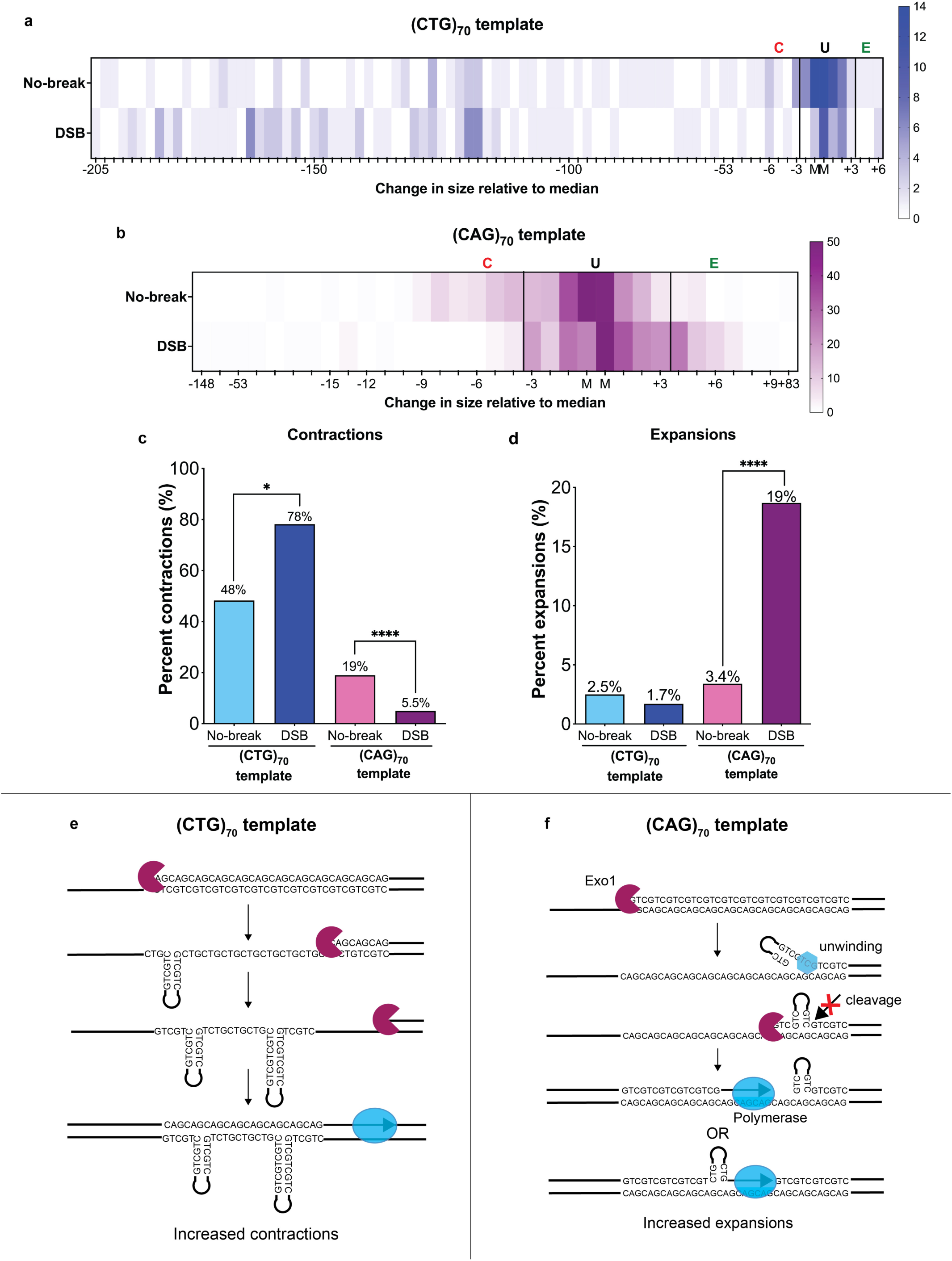
Gap filling during repair results in repeat instability. **a)** Heat map showing repeat length changes measured in the (CTG)_70_ template strain. M denotes median tract lengths of no-break condition. Region between black lines is within the error of the analysis and thus designated as unchanged (U). Expansions are denoted as (E), contractions denoted as (C). Tick marks denote change in sizes in bp relative to the median. Total number of PCR reactions represented: no-break n=120, break n=119. **b)** Heat map showing repeat length changes measured in the (CAG)_70_ template strain. Abbreviations are the same as in A. Total number of PCR reactions represented: no-break n=261, break n=273. **c)** Quantification of contractions. Contractions were determined by counting number of colonies below the unchanged thresholds determined in (A) and (B). Left: for the (CTG)_70_ template, contractions significantly increased in the DSB condition compared to the no-break condition, p=0.02. Right: for the (CAG)_70_ template, contractions significantly decreased in the DSB condition compared to the no-break condition, p=0.0001. Statistical analysis by Fisher’s exact test. **d)** Quantification of expansions. Expansions were determined by counting number of colonies above the unchanged thresholds determined in (A) and (B). Left: for the (CTG)_70_ template, expansion frequencies are not significantly different between the no-break and DSB conditions. Right: for the (CAG)_70_ template, expansions significantly increased with DSB induction compared to the no-break condition, p=0.0001. Statistical analysis by Fisher’s exact test. **e)** Model for repeat contractions during gap filling of the (CTG)_70_ template. Resection over the (CTG)_70_ template occurs unimpeded. The resulting ssDNA is left unprotected such that DNA hairpins can form. Polymerase fill-in bypasses the hairpins resulting in contraction of the repeat tract. **f)** Models for repeat expansions in the (CAG)_70_ template strain. During resection, helicase unwinding or strand displacement results in a ssDNA flap on the strand being resected that forms a small CTG hairpin which is resistant to endonuclease cleavage. Failure to resolve the hairpin on the resected strand results in incorporation of that sequence during the restoration of the dsDNA molecule. Alternatively, polymerase slippage through the repeat tract results in the addition of bases.

Expansions and contractions were defined as ≥3 bp above (E, expansion) or ≥3 bp below (C, contraction) the median determined in the no-break condition; regions between the lines are defined as unchanged (U) (Figure 3a, 3b). In the assay system used here, the repeat tract was inserted into the *ILV6* locus on chromosome III, 4 kb away from the ARS307 origin of replication which is to the right of the *HPH* gene (Figure 1a). Thus, in the (CTG)_70_ template strain, the CTG sequence is on the lagging template strand during replication. Consistent with previous work ^22–, 24^, we note a contraction frequency of 48% in the no-break condition. Interestingly, contraction frequencies significantly increase in the DSB condition to 78% (Fisher’s exact test, p=0.028) (Figure 3c). Separation on the fragment analyzer allowed for precise determination of how many repeat units were either gained or lost. In the (CTG)_70_ no-break condition, the size of the contractions was between 1 and 64 repeats (3 to 193 bp), with a median loss of 41 repeats (123 bp). This contraction size range is somewhat larger in the break condition, where colonies lost between 1 and 68 repeats (3 to 205 bp) and had a median loss of 48 repeats (144 bp). There is no significant break-dependent change in repeat expansions in the (CTG)_70_ template strain (no-break: 2.5%, break condition: 1.7%) (Figure 3d). Together, these data show that gap filling increases contraction frequency and large-scale contractions are favored when CTG is on the ssDNA template strand.

In contrast, instability in the (CAG)_70_ template strains shifted to larger (expanded) sizes after DSB repair compared to the no-break condition (Figure 3b). Interestingly, most of these expansion events are small and only add 1 repeat (3 bp) to the repeat tract; 88% of expansions are within 1 repeat of the expansion cutoff (Figure 3b). In the no-break condition, there is a 3.4% expansion frequency, whereas in the break condition, the expansion frequency significantly increases to 18.7%, a 5.5-fold increase (Fishers exact test, p=0.0001) (Figure 3d) showing that gap filling is driving small repeat expansions. The shift towards expansions also resulted in a shift away from contractions when (CAG)_70_ is the fill-in template (Figure 3c). The contraction frequency is 19.2% in the no-break condition and significantly decreases to 5.5% in the break condition, a 3.5-fold decrease (Fishers exact test, p=0.0001). The reduction in contractions supports that the no-break and break conditions are measuring instability due to different biological events, e.g. replication or repair of naturally occurring breaks within the repeat versus gap filling over a single-stranded repeat. To confirm that gap fill-in mediated instability was a function of the structure-forming ability of the repeat tract, we determined the instability of the scrambled control sequence (SFigure 3c). Though some instability exists for the scrambled control, expansion and contraction frequencies were not significantly different between the no-break and break conditions (expansions, no-break: 2.6%, break 2.1%; contractions, no-break: 2.6%, break 3.5%) (SFigure 3d).

The differences in expansion and contraction frequency between the (CAG)_70_ and (CTG)_70_ template conditions suggest the following model for the causes of CAG/CTG repeat instability during gap filling. In the (CTG)_70_ template strains, there’s impaired repair and increased contractions but resection occurs unimpeded. The lack of a resection defect could be explained by the fact that the CAG tract on the 5’ recessed end forms a less stable hairpin, so is more easily unwound and processed. On the other hand, the ssDNA exposed on the template strand by resection can form stable CTG DNA hairpins. Bypass of these hairpins during polymerase mediated fill-in could result in the frequent and large-scale contractions observed (Figure 3e). Alternatively, a break that occurs within the single-stranded CTG template after annealing of the U2 homologies (so that gap filling provides a CAG top strand) could be repaired such that out of register alignment results in a repeat contraction (Figure S3e). In the (CAG)_70_ template strains, resection is impaired suggesting a CTG hairpin forms on the 5’ end of the resected strand which is resistant to processing. To explain the increase in gap fill-in dependent expansions two models can be envisioned that could lead to small expansions. One possibility is that the hairpin on the resected strand remains unresolved and is incorporated by ligation after polymerase fill-in (Figure 3f). Another possibility is that expansions occur by polymerase slippage during gap filling (Figure 3f). These possibilities are not mutually exclusive: slippage may be further promoted by the presence of a hairpin on the 5’ flap which could impair polymerase release and ligation. We cannot exclude that other models for repeat length changes could exist. Regardless, these data show that the template of a repeat tract with respect to repair synthesis determines its stability in the genome and whether it will be more prone to expansions or contractions.

### Breaks at the single-stranded CTG tract explain the decreased viability in the CTG template strain

There is a dramatic decrease in viability in the (CTG)_70_ template strain (Figure 1c). We hypothesized that this may be due to fragility at the CTG repeat tract post DSB induction at the HO site, resulting in two breaks. The assay system we constructed had a second duplicated sequence, as the MX cassettes share homologies on either side of the *HPH* and *NAT* drug-resistance markers (Figure 4a, dark green boxes). If a break at the single-stranded CTG repeat tract occurred, the MX homologies could be a site of recombinational repair; however, this larger deletion would also delete the essential *NFS1* gene and be inviable. To determine if this event was occurring, we tested whether supplying the cell with a copy of *NFS1* would be sufficient to rescue the viability defect. Using genetic complementation of *NFS1* on a single copy plasmid, we found that complementing *NFS1* resulted in a significant increase in viability of the (CTG)_70_ template strain compared to the vector only control (Figure 4b). In the (CTG)_70_ + *NFS1* complementation experiments there were two populations of colonies. The first grew as expected and had an amplifiable CTG tract (large colonies; SFigure 4a). The second, which only appeared after *NFS1* complementation, grew poorly, and did not have an amplifiable repeat tract, suggesting these were the colonies in which the larger deletion had occurred (small colonies; SFigure 4a). As the small colonies were difficult to propagate, it is likely that while complementation with *NFS1* is sufficient for rescuing viability, loss of additional genes on the region between the HO site and the repeat tract predicted to be deleted, such as the RFC protein Dcc1, also impairs cellular fitness.

**Figure 4.**
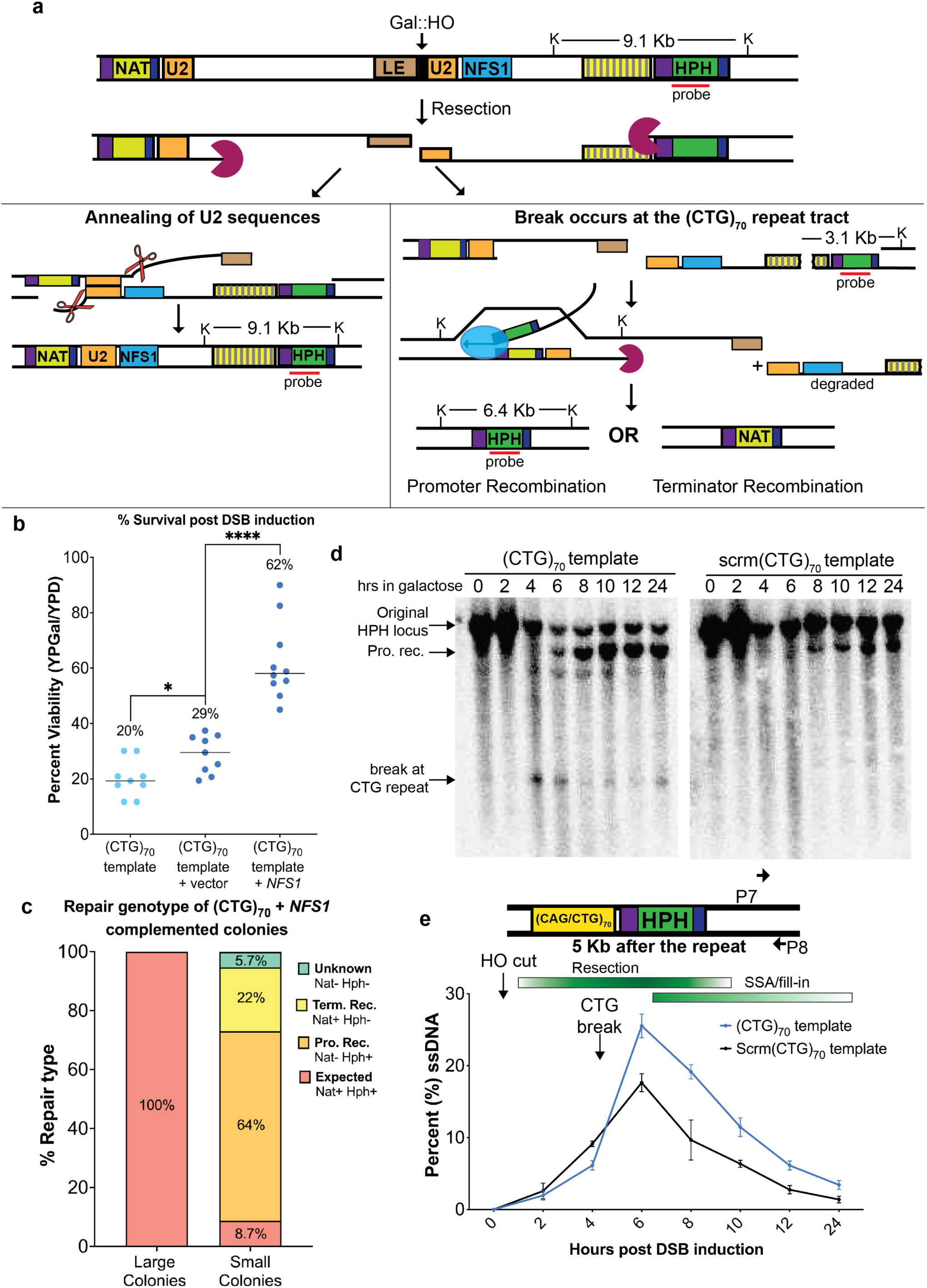
A second DNA break occurs at the (CTG)_70_ repeat and results in decreased viability. **a)** The model of the two-break hypothesis in the (CTG)_70_ template. Upon DSB induction, resection occurs on both sides of the break. *Left*, if repair occurs as expected, the two U2 regions of homology anneal, and error-prone gap filling occurs through the repeat tract leading to contractions. *Right*, if a second break occurs at the (CTG)_70_ repeat tract during resection, then repair occurs via BIR using the TEF promoter or TEF terminator sequences as homologies which are present in nearby marker genes. Recovered alternatively repaired colonies are either NAT+ or HPH+. The genomic region between the two breaks spans chromosome III position 92418-104619. The ssDNA fragment is subject to exonucleolytic degradation and loss of the essential gene *NFS1*, as well as the *DCC1*, *BUD3*, *YCL012C*, *GBP2*, and *SGF29* genes. *HPH* probe and relevant *Kpn*I sites and expected sizes using the *HPH* probe are marked. The expected size of the *Kpn*I-digested fragment of the original *HPH* locus is 9.1 kb and recombination products between the TEF promoter (Pro. Rec) produces a band of ∼6.4 kb. If a break occurs at the CTG repeat, a band ∼3.1 kb is expected. **b)** Percent viability of (CTG)_70_ template (n=9), (CTG)_70_ template + vector (n=9) and (CTG)_70_ template+*NFS1* (n=10) are shown. (CTG)_70_ no vector to (CTG)_70_+vector is p=0.0134 and to (CTG)_70_+*NFS1* is p<0.0001 by Student’s t-test. **c)** Genetic typing of large (n=69) and small (n=115) repaired colonies in the (CTG)_70_ template strain that was complemented with *NFS1*. **d)** Kinetic Southern blots of KpnI digested DNA were stripped and probed with a fragment to the *HPH* locus. Representative Southern shown; number of replicates: scrm(CTG)_70_ (n=2) and (CTG)_70_ (n=4). **e)** Resection and gap filling kinetics of a region 5 kb after the repeat was determined for scrm(CTG)_70_ (n=2) and (CTG)_70_ (n=4) strains as described in Figure 2b.

This assay system can utilize both SSA and BIR repair pathways for repair^21^. If recombinational repair mechanisms like BIR were occurring between the MX cassettes, there are two possible homologies that are present in the *NAT* and *HPH* marker genes that could be used: the TEF promoter (344 bp) or the TEF terminator (198 bp). Recombination between the TEF promoters would result in the loss of the *NAT* marker gene, while recombination between the TEF terminators would result in the loss of the *HPH* marker gene. To test whether alternative recombination between markers could explain the loss in viability, we replica-plated all colonies on media containing either nourseothricin or hygromycin. As expected, 100% of large colonies contained both the *NAT* and *HPH* markers (Figure 4c) indicative of repair using the expected U2 homology (Figure 4a, left pathway). However, small colonies only contained both markers 8.7% of the time (Figure 4c), consistent with alternative repair occurring most of the time (Figure 4a, right pathway). Small colonies retained the *HPH* marker 64% of the time and the *NAT* marker 22% of the time, consistent with more recombination occurring at the TEF promoters that share more homology.

If the (CTG)_70_ ssDNA forms stable hairpins, they could be a substrate for nucleolytic cleavage resulting in a second break^28^. To see if a second break could be detected, the Southern blots monitoring SSA from Fig 2A were re-hybridized with a probe to the *HPH* gene 600 bp downstream from the repeat. We observed a unique band in the (CTG)_70_ template strain that corresponds to the size expected if the break occurred within the (CTG)_70_ repeat tract (Figure 4d). This break was not observed for the CAG template or no repeat strains (Figure S4b). In addition, a band corresponding to the size expected for recombination between the TEF promoters was observed in the CTG repeat strain (Figure 4d). Recombination between TEF promoters is also observed in the Scrm(CTG)70, (CAG)_70_ and no repeats strains though at a reduced level compared to the (CTG)_70_ template strains (Figure 4d; Figure S4b). The break at the (CTG)_70_ template strain appears at hour 4, suggesting the repeat tract breaks when the DNA in this region has become single-stranded (Figure 4d; Figure 2b) and that repeat tract breakage is promoting the increase in TEF promoter recombination. There seems to be some portion of cells that cannot repair using the alternative homologies within the time course, resulting in the persistence of the CTG repeat-specific band beyond 24 hours. Further, a portion (5.7%) of small, complemented colonies lose both the *HPH* and *NAT* markers (Figure 4c), suggesting other recombination events are likely occurring. Together, these results indicate that breakage occurs at the (CTG)_70_ tract when it becomes single-stranded, resulting in alternate repair events that cause a loss of viability.

We considered whether the second break and alternative repair between promoters could help explain the increased ssDNA profile observed during resection across the (CTG)_70_ template (Figure 2b). BIR occurs via conservative DNA synthesis and has asynchronous replication of the leading and lagging strands^29^. If BIR were occurring at the site of the TEF promoter and synthesizing to the telomere end via D-loop bubble migration, then persistent, increased ssDNA would be observed at sites, like in the *HPH* gene (Figure 2b), that had yet to be filled in (Figure S4c). Consistent with delayed gap filling due to BIR, increased, persistent ssDNA was detected at a location ∼5 kb after the (CTG)_70_ template repeat tract compared to the scrm(CTG)_70_ control (Figure 4e, P7&8).

We also tested whether a possible reason for the increased BIR between the *HPH* and *NAT* loci was from CTG breakage events that had occurred before resection exposes the U2 homology. Supportive of this hypothesis, DNA 2.3 kb before the U2 homology (i.e., ∼23 kb from the HO-induced DSB) is minimally single-stranded at hour 4 when the break at the (CTG)_70_ repeat tract occurs (SFigure 4d). Interestingly, at this location, the resection and fill-in kinetics appear to be delayed once the break at the repeat tract occurs, suggesting the possibility that the DNA substrate accessibility to exonucleases changed because of the ongoing BIR nearby. In summary, exposure of ssDNA by resection over the structure forming CTG tract creates a highly fragile site, resulting in altered repair, large-scale deletions of neighboring genes, and cell death.

### Regulation of resection by Rad9 increases repair kinetics to rescue viability and repeat-induced contractions

We next wanted to determine what genetic factors could impact repair efficiency during gap filling of a (CTG)_70_ template. Rad9 is an evolutionarily conserved DNA damage checkpoint protein that also has functions in restricting resection^8^. Loss of Rad9 results in increased recruitment of RPA, Rad51, and Rad52 on resected DNA around a DSB^12^. Intriguingly, deletion of *RAD9* almost completely rescued the loss in viability in the (CTG)_70_ template strain, but had no impact on viability in the scrm(CTG)_70_ strain (Figure 5a). In addition, deletion of *RAD9* showed a significant reduction in the frequency of repeat contractions during gap filling across the CTG tract (Figure 5b). It was previously established that resection speed in *rad9*Δ mutants is twice as fast as wildtype^10^. To test the role of Rad9 in resection in the presence of a structure-forming repeat, ssDNA levels were measured over the time course, monitoring a site after the repeat tract (primers P3 & P4). The resection kinetics were altered in the *rad9*Δ mutant, showing maximal ssDNA 2-4 hours earlier than in wildtype strains, and this was independent of the presence of the repeat tract (Figure 5c, SFigure 5a). Unlike in the wildtype (CTG)_70_ template strain, the *rad9*Δ mutant does not have persistent ssDNA at later time points (Figure 5c). Consistent with faster resection and filling in, there was earlier appearance of the SSA product in the *rad9*Δ mutant compared to wildtype (SFigure 5b, compare to Fig 2a). The increased speed of resection, repair, and fill-in in the *rad9*Δ mutant compared to wildtype could explain the rescue in (CTG)_70_ strain viability. Indeed, CTG repeat breakage could not be detected in the *rad9*Δ mutant when Southern blots monitoring SSA were re-probed to the *HPH* locus (Figure 5D), indicating that loss of Rad9 results in fewer breaks at the repeat locus. To test whether faster recruitment of RPA and Rad51 could help protect ssDNA from hairpin formation and nucleolytic cleavage, we employed ChIP analysis across the full time-course. Deletion of *RAD9* resulted in earlier recruitment of RPA and Rad51 at the (CTG)_70_ tract compared to the wildtype (Figure 5e & f, see hour 2), which is consistent with the earlier peak of ssDNA accumulation at 2 hours (Figure 5c). Correspondingly, RPA is removed more quickly in the *rad9*Δ mutant, disappearing by 6 hours, whereas RPA is maximal at 6 hours in wildtype cells. Rad51 enrichment was somewhat lower but also followed the faster repair kinetics of repair in the *rad9*Δ mutant compared to wildtype. Taken together, as resection and repair are faster but there is not increased RPA and Rad51 in the *rad9*Δ mutant, this data suggests that the rescue in (CTG)_70_ contractions and viability in the *rad9*Δ mutant is due to faster repair kinetics which results in less opportunity for CTG hairpin formation and tract breakage.

**Figure 5.**
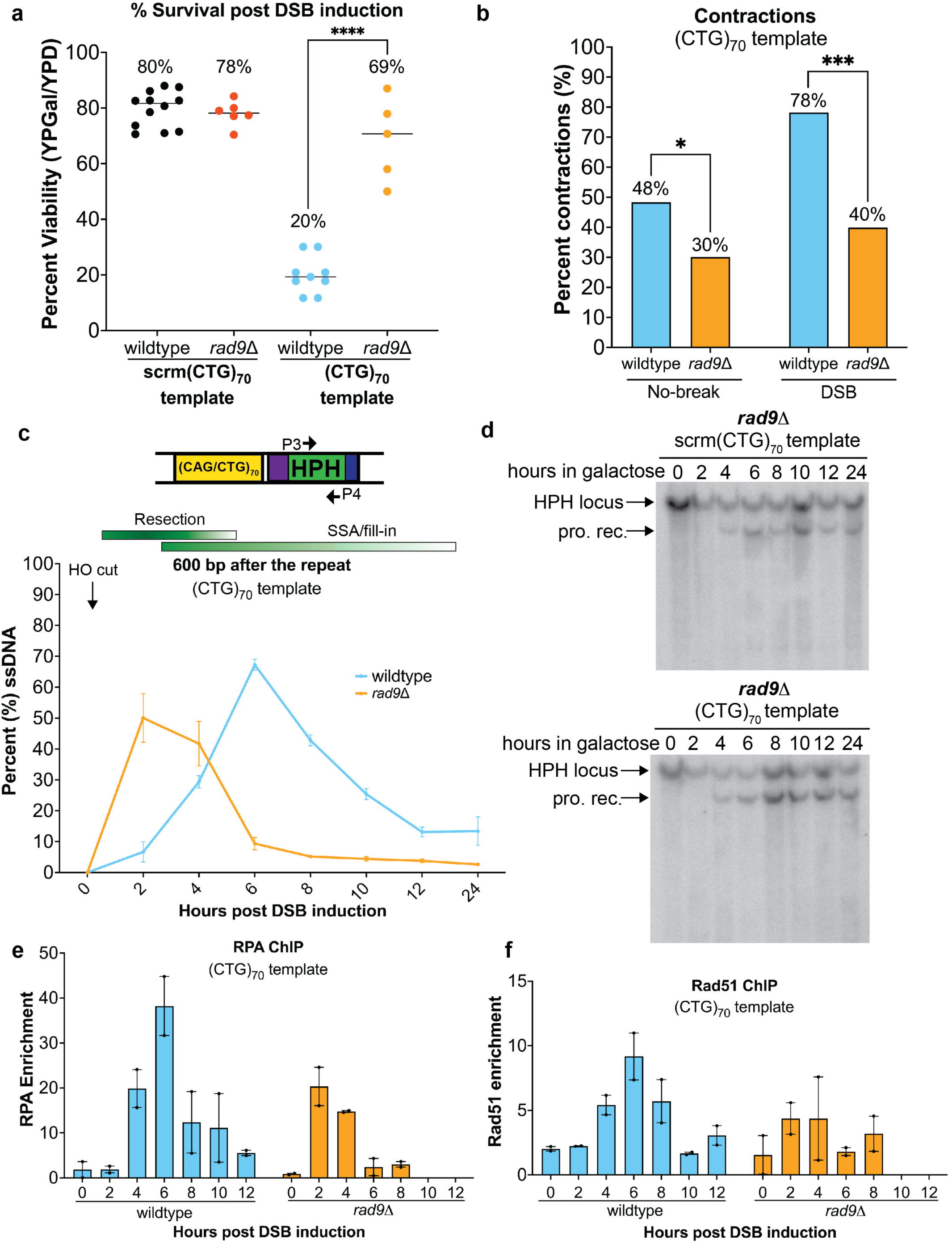
Deletion of *RAD9* rescues the decreased viability and gap fill-in mediated (CTG)_70_ contractions. **a)** For the scrm(CTG)_70_ template strain, percent viability of the *rad9*Δmutant (n=6) is unchanged compared to wildtype (n=12). For the (CTG)_70_ template strain, percent viability of the *rad9*Δ mutant (n=5) is significantly increased compared to wildtype (n=9) (p=0.0002). Statistics determined using Students t-test. **b)** In the (CTG)_70_ template strain, the contraction frequency in the no-break condition decreased in the *rad9*Δ mutant, p=0.04. In the DSB condition, the contraction frequency decreased more significantly in the *rad9*Δ mutant, p=0.001. Statistical analysis by Fisher’s exact test. **c)** Percent ssDNA 600 bp after the repeat locus was determined after DSB induction as in Figure 2b; wildtype n=4, *rad9*Δ n=3. **d)** Kinetic Southern blots were stripped and probed with a fragment to the *HPH* locus. Representative Southern shown; number of replicates: scrm(CTG)_70_ (n=3) and (CTG)_70_ (n=3). Enrichment of **e)** RPA and **f)** Rad51 occurs earlier near the (CTG)_70_ repeat tract in the *rad9*Δ mutant following DSB induction compared to wildtype. Independent biological replicates for wildtype (n=2) and *rad9*Δ (n=2). Enrichment adjacent to the (CTG)_70_ repeat was determined using primers P3 & P4 and calculated using absolute quantity and normalized to *ACT1*. Graph shows mean ± SEM.

### Rad51 protects the single-stranded (CTG)_70_ repeat from fragility and contractions

Due to the extensive amount of ssDNA produced during this SSA repair assay and its importance in determining repair outcome, we tested the role of Rad51, which binds ssDNA to promote strand exchange. Rad51 is not required for repair via SSA^15^, but it has been shown that the parent assay system can repair via either SSA or Rad51-dependent BIR, with similar kinetics^21^. Indeed, Rad51 is recruited to the ssDNA during resection in both the scrm(CTG)_70_ and (CTG)_70_ template strains (SFigure 2). Deletion of *RAD51* in the (CTG)_70_ template strain resulted in a significant decrease in viability (Figure 6a). Deletion of *RAD51* in the scrm(CTG)_70_ template also had a decrease in viability, though less dramatic (Figure 6a). Lack of Rad51 could result in less protection of the repeat on the single-stranded template leading to secondary structures that are targets of nucleolytic cleavage or contraction intermediates (Figure 4a).

**Figure 6.**
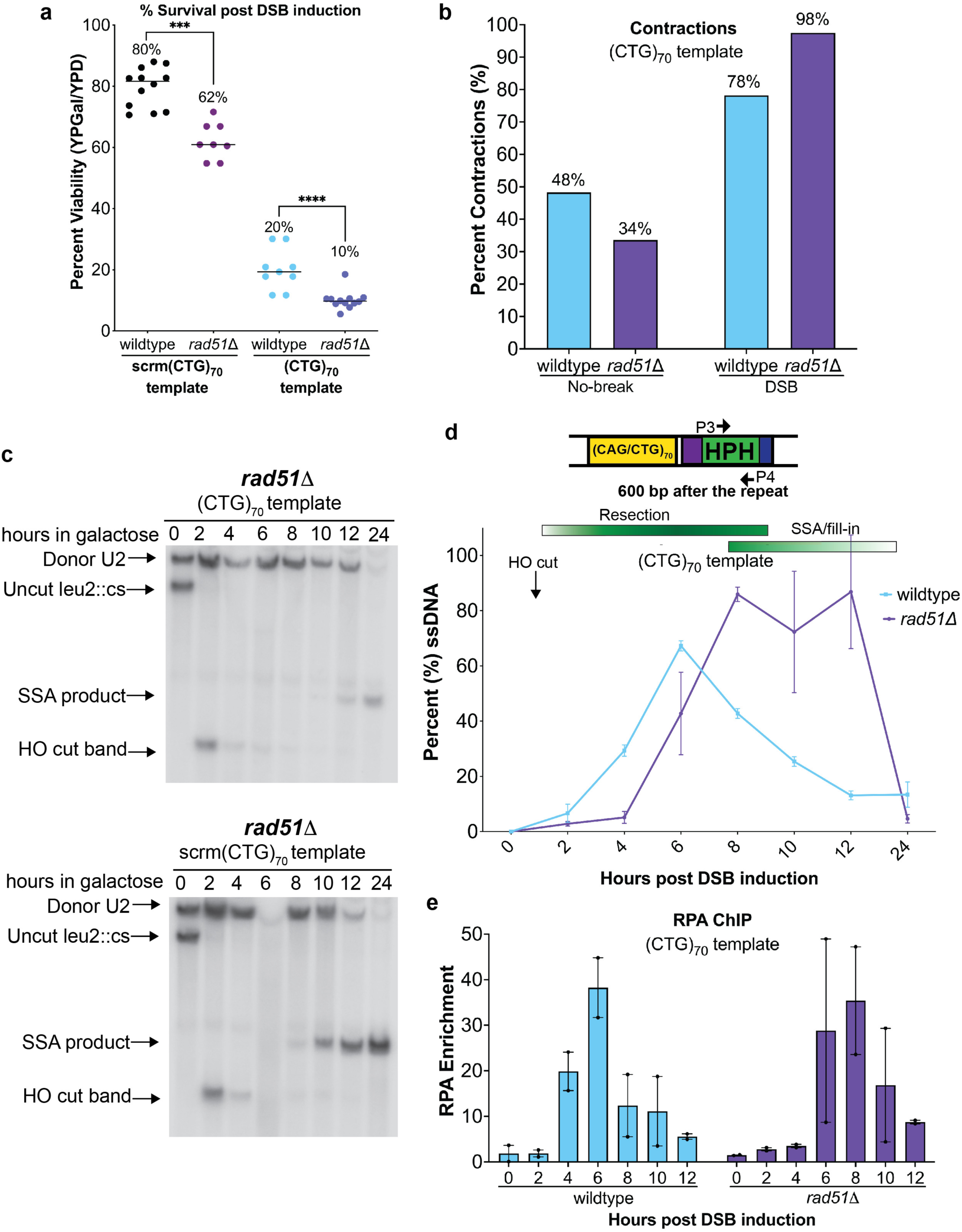
Deletion of *RAD51* impairs repair and results in increased contractions during gap filling. **a)** For the scrm(CTG)_70_ template strain, percent viability of the *rad51*Δ mutant (n=8) is significantly decreased compared to wildtype (n=12) (p=0.002). For the (CTG)_70_ template strain, percent viability of the *rad51*Δ mutant (n=12) is significantly decreased compared to wildtype (n=9) (p=0.0002). Statistics determined using Students t-test. **b)** For the (CTG)_70_ template no-break condition, *rad51*Δ mutants had decreased contractions compared to wildtype (p=0.15). In the DSB condition, *rad51*Δ mutants had increased contractions compared to wildtype (p=0.26). Statistical analysis by Fisher’s exact test. **c)** Kinetics of repair for the scrm(CTG)_70_ (n=2) and (CTG)_70_ (n=2) template strains in *rad51*Δ mutants. Representative Southern shown; probe is to a portion of the *LEU2* gene (probe location noted in Figure 1a). **d)** Resection and fill-in kinetics for the (CTG)_70_ template in wildtype (n=4) and *rad51*Δ (n=2) strains after DSB induction. Percent ssDNA after the repeat locus was determined as in Figure 2b. **e)** Enrichment of RPA near the (CTG)_70_ repeat tract in wildtype and *rad51*Δ mutant following DSB induction. Independent biological replicates for wildtype (n=2) and *rad51*Δ (n=2). Enrichment adjacent to the (CTG)_70_ repeat was determined using P3 & P4 and calculated using absolute quantity and normalized to *ACT1*. Graph shows mean ± SEM.

Consistent with this idea, loss of Rad51 increased the frequency of CTG repeat tract contractions during gap filling to nearly 100% (Figure 6b).

To better determine whether the addition of a repeat tract changes repair kinetics in the *rad51*Δ mutant, we followed the time course of the SSA reaction by Southern blot (Figure 6c). Consistent with previous work^21^, the appearance of the repair product was delayed by ∼4 hours in the scrm(CTG)_70_ strain (Figure 6c), first appearing around 8 hours instead of 4 hours and then increasing slowly from 10-24 hours. The presence of the (CTG)_70_ template led to a dramatic delay, with repair products not appearing until 10-12 hours post incision in the absence of Rad51 (Figure 6c). Breaks that occur at the CTG repeat tract appear when the DNA first becomes single-stranded and persist through hour 8 (SFigure 6a). Interestingly, alternative recombination between the TEF promoters is reduced and delayed compared to wildtype (Compare SFigure 6a to Figure 4d) suggesting that Rad51-dependent repair is driving promoter recombination.

Given that Rad51 seems to play a protective role at the CTG repeat tract, we measured the kinetics of generating ssDNA after the repeat locus in the *rad51*Δ mutant. Interestingly, *rad51*Δ mutants had a 2-4 hour delay in resection through the repeat locus in the scrm(CTG)_70_ and (CTG)_70_ template strains (Figure 6d, SFigure 6b). The delayed resection in the *rad51*Δ mutant is accompanied by delayed enrichment of RPA 600 bp after the repeat (Figure 6e). The delay in ssDNA accumulation is consistent with the delay in repair in these strains. Strikingly, the region after the (CTG)_70_ template is maximally single-stranded through the 12-hour timepoint in the *rad51*Δ mutant compared the wildtype (Figure 6d). There is also increased ssDNA in the *rad51*Δ mutant in the scrm(CTG)_70_ strain though it does not persist to the same degree (SFigure 6b). Thus, a deficiency in Rad51 binding causes ssDNA persistence, which is exacerbated when a structure-forming CTG repeat is on the exposed single-stranded template strand. The loss of the competing BIR pathway likely explains the delayed appearance of the repair product in the absence of Rad51, the persistent ssDNA and the additional loss of viability in this mutant. Taken together, these data reveal a dual role for Rad51, first in protecting repeat tracts from forming DNA structures during resection which prevents contractions and fragility, and then by providing BIR as a repair pathway choice. The repair via BIR may be especially important when there are multiple breaks due to a fragile sequence.

## Discussion

Previous work had shown that CAG repeat expansions and contractions can occur during HR repair, but it was not clear which steps of HR were involved. In this study, we developed an assay system to test the stability of expanded CAG/CTG trinucleotide repeats specifically during the gap filling step of HR. Our data show that both resection and subsequent DNA polymerization during gap repair are mutagenic steps of HR when a structure-forming CAG/CTG tract is present. The identity of the repeat tract in relation to the resected strand results in very different phenotypes. The (CAG)_70_ template, with CAG on the ssDNA template strand and CTG on the 5’ recessed end, has resection defects and is prone to gap fill-in mediated expansions. When the repair template strand has a (CTG)_70_ repeat there are two possible repair outcomes (Figure 4a). First, if gap filling occurs as expected this frequently results in large scale contraction of the CTG repeat tract. The surprising second outcome is a repeat tract-dependent single-strand break that occurs after resection, leading to altered repair and cell death. We discovered that Rad9’s normal function in slowing down resection can have a detrimental effect during repair of gaps containing structure-forming repeats, presumably by allowing opportunity for structure formation on the exposed single-stranded template. Finally, we discovered that Rad51’s presence protects against repeat contractions and provides an opportunity for a BIR-mediated repair pathway when there is repeat associated fragility.

In the situation where the gap repair template strand was an expanded (CAG)_n_ tract (CAG template) and the more stable CTG structure could form on the 5’ resected end, there were decreased levels of resection beyond the repeat and an increase in expansions.

Supportive of the possibility that non-B form structures can be a barrier to resection, it has been shown that resection is impeded and repair choice is altered in the presence of a stabilized G-quadruplex on the resected strand^7^. Because resection of this strand to the distance of the TNR is not imperative for repair, viability and repair kinetics were unaltered. However, repeat stability was altered so that gap fill-in outcomes favored CAG repeat expansion. The decreased resection coupled with the increase in expansions provides a simple model for repeat expansion due to incorporation of an unprocessed structure on the 5’ flap during the final ligation step of gap filling (Figure 3f). What is surprising about these expansions is how relatively small they are, as most of them are an addition of only 1 or 2 repeat units (3-6 bp). These small expansions stand in contrast to large-scale CAG expansions that were shown to occur during a BIR-like process dependent on Rad52 and Pol32^31, 32^. It suggests that the hairpins formed on the resected strand are relatively small. It is also possible that the DNA structure impairing resection is resolved and that the expansion event is due to polymerase slippage, or that the two processes are linked (Figure 3f). These small-scale expansions could be like those that occur during gap repair in non-dividing human cells, such as neurons, that undergo stepwise somatic expansions which advance disease onset in trinucleotide repeat expansion disorders such as Huntington’s disease^6^.

In the situation where the gap repair template strand was an expanded (CTG)_n_ tract (CTG template), contractions predominated and there was a loss in viability. In cells that repaired as expected (using the U2 homology), the CTG repeat likely forms several hairpins which are bypassed during gap filling leading to a variety of large-scale contractions (Figure 3e, 7a). Alternatively, DNA breaks in the single-stranded CTG repeat tract could result in contraction via out of register alignment (SFigure 3e). This second model for repeat contractions is similar to that proposed for Cas9-induced breaks at an expanded CAG/CTG repeat which repair via end joining or SSA^26^ except that the break occurs when the repeat tract is single-stranded, rather than it being an induced DSB. During gap repair of the (CTG)_70_ template strain there was also a surprising secondary event: a frequent resection-dependent break within the CTG repeat that led to the formation of a large, lethal deletion. In this assay system, the large gap (up to 25 kb) creates a long stretch of ssDNA and a significant need for ssDNA protection during repair. Our data reveal that this situation is particularly dangerous for the cell as the ssDNA is prone to breakage, and this leads to large deletions and loss of vital genetic material (Figure 7a). Similar events could be behind large deletions that often occur at fragile sites in the human genome^33^. The significant difference between having the CAG or CTG strand on the exposed template indicates that structure-forming potential is a key determinant of whether a ssDNA region results in chromosome fragility.

**Figure 7:**
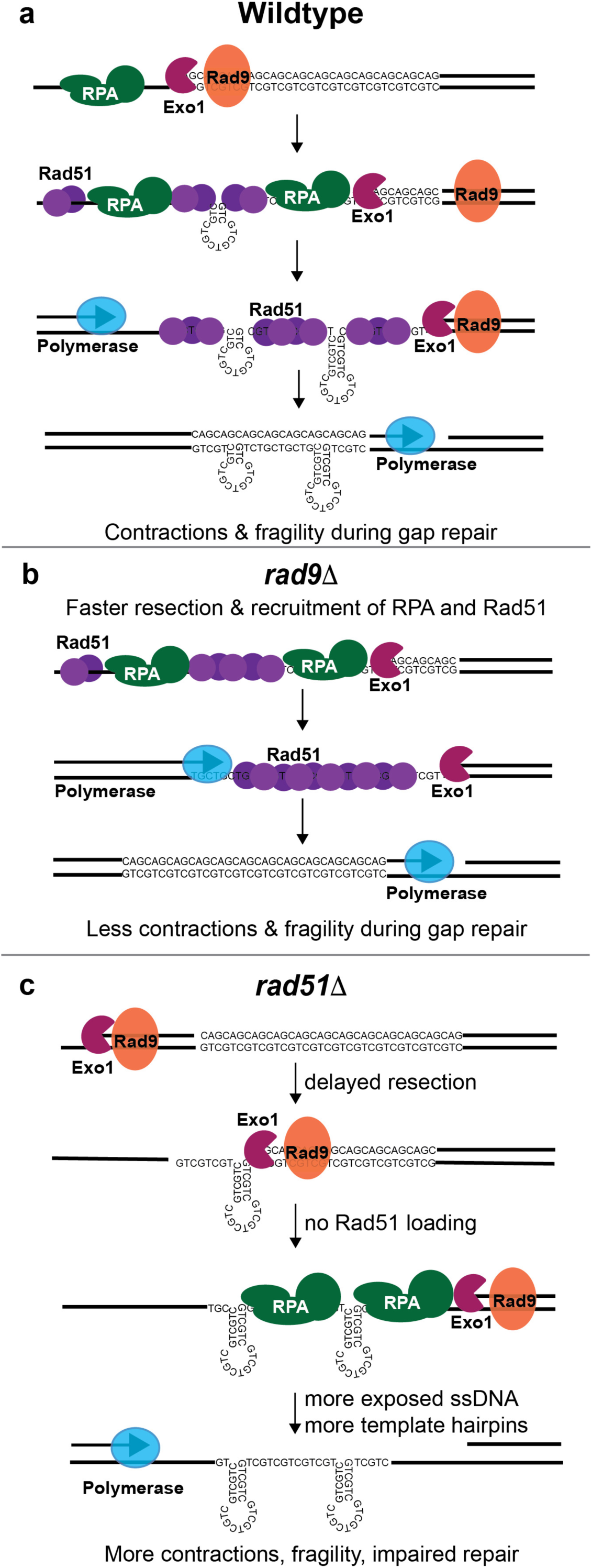
A model for repeat instability during resection and gap filling at a CTG repeat. **a)** In wildtype cells, as resection occurs, Rad9 controls the recruitment of RPA and Rad51 to ssDNA. Delayed binding of RPA or Rad51 results in hairpin formation. Contractions are a result of hairpin bypass during polymerase mediated gap filling (shown) or by SSA/end-joining events within the broken CTG repeat region (SFig3e). In addition, exposed ssDNA results in increased breakage and alternative repair. **b)** In *rad9*Δ mutants, there no delayed recruitment of RPA and Rad51 to the freshly resected ssDNA. This prevents hairpin formation and breakage at the repeat tract resulting in increased viability and decreased contractions. **c**) In *rad51*Δ, initiation of resection is delayed. Absence of Rad51 and slowed gap filling both allow for increased DNA structure formation and contractions. Breaks that occur at the CTG repeat tract cannot repair via BIR leading to increased cell death.

One of the more surprising findings was that repeat instability and fragility of the (CTG)_70_ template was reduced in the absence of Rad9, the *S. cerevisiae* ortholog of 53BP1. We confirmed that resection is faster in the *rad9*Δ mutant and there is earlier recruitment of RPA and Rad51, suggesting a model where faster recruitment of RPA and Rad51 help guard against hairpin formation. One of the major roles of RPA is to prevent DNA secondary structure formation. RPA is a trimeric complex that normally coats ∼25 nt of ssDNA^34^ while Rad51 is much smaller and coats ∼3 bp of ssDNA^35^. Since we see RPA and Rad51 accumulate as the DNA is resected, one possibility is that in wildtype cells a brief lag between resection and RPA or Rad51 loading normally occurs^36^ and is sufficient to allow formation of small hairpins that form as resection proceeds (Figure 7a). Loss of Rad9 may eliminate the lag between resection and RPA or Rad51 loading, resulting in fewer DNA hairpins and a more stable repeat tract (Figure 7b). An alternate hypothesis is that Rad9 normally delays gap filling through interactions with polymerases and other accessory factors. In mammalian cells, it has been suggested that unrestricted resection may not be the only model for the increased ssDNA in the absence of 53BP1, rather it could also be due to impaired gap filling^9, 37^. Recent work has shown that 53BP1 facilitates gap filling by recruitment of Shieldin and subsequently the CST complex and Pol- alpha primase^38, 39^. However, there may be differences between the functions of Rad9 and 53BP1 as our data shows that gap filling occurs without delay in the absence of Rad9 and any role Rad9 may have in recruitment of gap filling factors is not essential for completion of fill-in synthesis.

We’ve identified novel roles for Rad51 in long range resection and in protection of repetitive ssDNA. Loss of *RAD51* resulted in increased gap repair mediated contractions (Figure 6b) and inviability (Figure 6a) when (CTG)_70_ was the fill-in template. The increase in contractions suggests that Rad51 may have a protective role in gap repair by binding ssDNA and preventing secondary structure formation (Figure 7c). Interestingly, *rad51*Δ mutants have a slower rate of 5’ to 3’ resection (Figure 6d, SFigure 6b) and this is mirrored by delayed RPA recruitment (Figure 6e), even in strains without a repeat tract. However once resection starts, *rad51*Δ mutants accumulate more ssDNA that persists for longer compared to wildtype. In our assay, Rad51-dependent BIR is one mechanism for repair^21^. Our data suggest that if a second break occurs at the CTG repeat, BIR using the MX homologies becomes the preferred mechanism for repair (Figure 4a, SFigure 4c). Elimination of the BIR pathway by deletion of *RAD51* leaves cells with a break that must repair via SSA using unintended homologous sequences and this is a major cause for the delayed repair and loss in viability in the *rad51*Δ mutant (Figure 7c).

Gap filling is not a process unique to HR; mismatch, base excision and nucleotide excision repair pathways all involve gap filling post-removal of a lesion, and repeat instability in somatic tissues affected in TNR diseases has been attributed to these pathways^25^. In addition, cells that repair DSBs using MMEJ exhibit increased mutation levels, presumably due to extensive resection of DSBs and mutagenic gap filling^40^. Further, during BIR there is an accumulation of ssDNA that will be filled in and this is a mutagenic event^29^. Recent work has shown that an internal telomeric repeat (ITS) impedes BIR leading to fragility and ITS instability^41^. Thus, our findings could also apply to other repair pathways that involve a gap filling step. Furthermore, TNR expansions accrue in non-dividing cells during aging^25^. As there are increased DSBs in neuronal tissues with age^42^, it is possible that repair of neuronal DSBs near sites of TNRs could lead to gap filling mediated instability, or alternative repair that could result in cell death.

Taken together we show that both resection and gap repair through a repeat tract can be mutagenic, creating expansions, contractions, or chromosome breakage depending on the location of the structure-forming sequence. We showed that structures on the single-stranded template strand cause either repeat contractions or chromosome breaks, whereas structures on the 5’ recessed strand interfere with resection and lead to small repeat expansions. More broadly, our results indicate that the success and accuracy of repair is influenced by the sequence context where repair is occurring and illustrate the danger of exposing ssDNA within repetitive sequences during gap repair

## Methods

### Yeast strains

All strains are derivatives of YMV80^15^ which was derived from S288C. (CAG)_70_, (CTG)_70_ and scrm(CTG)_70_ repeat tracts were integrated at the *ILV6* locus on chromosome V. Proper integration of the repeat tract was confirmed via Southern blotting and expected tract length was confirmed via PCR. Gene deletions were generated by one step gene replacement with a marker gene. All strains are listed in Table 1.

### Viability assay

Tract length was confirmed via colony PCR using primers listed in Table 1. Size was determined by either electrophoretic analysis using a fragment analyzer or 2% metaphor agarose. Colonies with confirmed tract length were inoculated into 2 mL YP+Lactate (pH 5.5) and grown for 2-3 divisions (16-18 hours). Cultures were appropriately diluted and plated on YPD and YP+ 2% Galactose (YPGal) in duplicate. Plates were incubated at 30°C for 2-3 days. Colonies were counted and percent viability was obtained by dividing the number of colonies of galactose by the number of colonies on glucose multiplied by 100. See Table 2 for individual assay values.

### SSA Repair Southern Blotting

Time course collection was adapted from^15^. Colonies of the correct tract length were inoculated into 10 mL YP+Lactate pH 5.5 for 24 hours. Cultures were diluted into 400 mL YP+Lactate pH 5.5 and grown for approximately 12-14 hours. Prior to galactose addition, cells went through approximately 6-7 cell divisions and the final cell number prior to addition of galactose was approximately 7 x 10^6^ cells per milliliter. Galactose was added to a final concentration of 2% and samples were taken at the indicated times. DNA was prepped via phenol-chloroform extraction and normalized via Qubit (dsDNA BR Assay Kit, Cat# Q32853; Invitrogen). DNA was digested with KpnI, separated on a 0.8% agarose gel, blotted, and probed with a fragment within the *LEU2* gene or *HPH* marker. Bands were visualized with a Typhoon phosphorimager (GE Biosciences).

### Resection/Gap fill-in assay

Adapted from^18^. DNA from the kinetic time courses was normalized using a Qubit. For each digest, 150 ng of DNA was digested with EcoRI or mock treatment overnight at 37°C such that the final concentration was 2 ng/ul. Each quantitative PCR (qPCR) reaction had 20 ng DNA and was run in duplicate. Determination of percent resected was done using %ssDNA= [100/[(1+2^ΔΔCt^)/2]/*f*] where ΔΔCt= ΔCt_digested_ - ΔCt_mock_ and *f* is HO cutting efficiency. HO cutting efficiency (*f*) was obtained using densitometric analysis using Imagequant. HO cutting efficiency was calculated as *f*=Cut value/ (Cut value+Uncut value).

Each resection assay has a paired Kinetic southern blot to monitor repair efficiency and was used to determine cutting efficiency. See Table 3 for individual values of each experimental replicate and primer set tested.

### CAG/CTG repeat analysis

For tract length analysis, resulting colonies from the viability analysis were used. Colony PCR was performed on 22-24 daughter colonies from both the paired YPD and YPGal plates using primers that span the repeat tract (Figure 1A). To eliminate variation, PCR and subsequent electrophoretic separation of repeat amplicons of daughter colonies from the YPD and YPGal conditions were done at the same time. For primers used see Table 1. For size analysis, PCR amplicons were sized on a fragment analyzer (Model# 5200; Agilent) using 600mer DNA separation gel (Cat# NDF-915-0275; Agilent) compared to a 1 bp & 6000 bp marker (Cat# FA-MRK915F-0003; Agilent) and a 100bp DNA plus ladder (Cat# FS- SLR915-0001; Agilent). For the *rad9*Δ and *rad51*Δ mutants, PCR amplicons were separated using standard gel electrophoresis on 2% metaphor agarose and sized in comparison to a 100 bp Hyperladder (Cat# BIO-33056; Bioline).

### Fragment Analyzer analysis

In order to determine the repeat size distributions of the no- break and break conditions, tract lengths (in bp) determined by the fragment analyzer were plotted versus the number of times the fragment analyzer made the same size determination. The median tract length in the no-break condition was mathematically determined. If the median was between two whole numbers, both whole numbers were designated as the median to determine expansion and contraction cut offs. The PCR amplicon (Figure 1A, Primers P1 & P2) encompasses 210 bp of the repeat tract and 150 bp of non-repetitive sequence. Due to the large spread of contractions, tract lengths below 300 bp were not used in the determination of the median in the (CTG)_70_ template. Expansions or contractions were any size called +/- 3 bp (one repeat unit) from the calculated median. Statistical significance was determined using Fisher’s exact test. See Table 4 for contraction and expansion frequencies for all strains listed. See Table 5 to see the number of times each size was called by the fragment analyzer.

### RPA and Rad51 Chromatin Immunoprecipitation

Time course was as described for the kinetic Southern blots. Samples for RPA and Rad51 ChIP were taken simultaneously from the same cultures. Cultures were crosslinked in 1% formaldehyde for 20 minutes and quenched with glycine (0.125 M final concentration) for 5 minutes. Immunoprecipitation was performed by incubating normalized samples with Protein G Dynabeads (Thermo Fisher, 10004D) pre- conjugated with a-RPA (2 uG; PA5-34905, Thermo Fisher Scientific) or a-Rad51 (3 uG; Agrisera AS07 214) for 2 hours at 4°C. Whole chromatin input and immunoprecipitated samples were subject to qPCR using primers 600 bp after the repeat locus (see Table 1) or *ACT1* as a control. Enrichment was determined using absolute quantity. See Table 6 for individual values of each experimental replicate.

## Acknowledgments

Research reported in this publication was supported by an American Cancer Society–Ellison Foundation Postdoctoral Fellowship PF-18-125-10-DMC to EJP and the National Institute of General Medical Sciences of the National Institutes of Health under Award Number R01GM122880 to CHF, R35 GM144215 to CHF, P01GM105473 to JEH and CHF, and R35 GM127029 to JEH. The content is solely the responsibility of the authors and does not necessarily represent the official views of the National Institutes of Health.

## Author Contributions

Experimental Design: EJP and CHF; Data accumulation: EJP, IDP and MDLR; Writing: EJP, JEH, and CHF.

## Tables

Table 1: Strains, Plasmids, Primers used

Table 2: Percent viability

Table 3: Listed experimental values for resection assays for all loci

Table 4: Instability frequency

Table 5: Fragment analyzer sizes called

Table 6: Enrichment of RPA and Rad51 by Chromatin Immunoprecipitation

