## Supplemental Figures 1-6, Supplemental Tables 1-6 for "Structure-forming CAG/CTG repeats interfere with gap repair to cause repeat expansions and chromosome breaks"

### Supplemental Figure 1

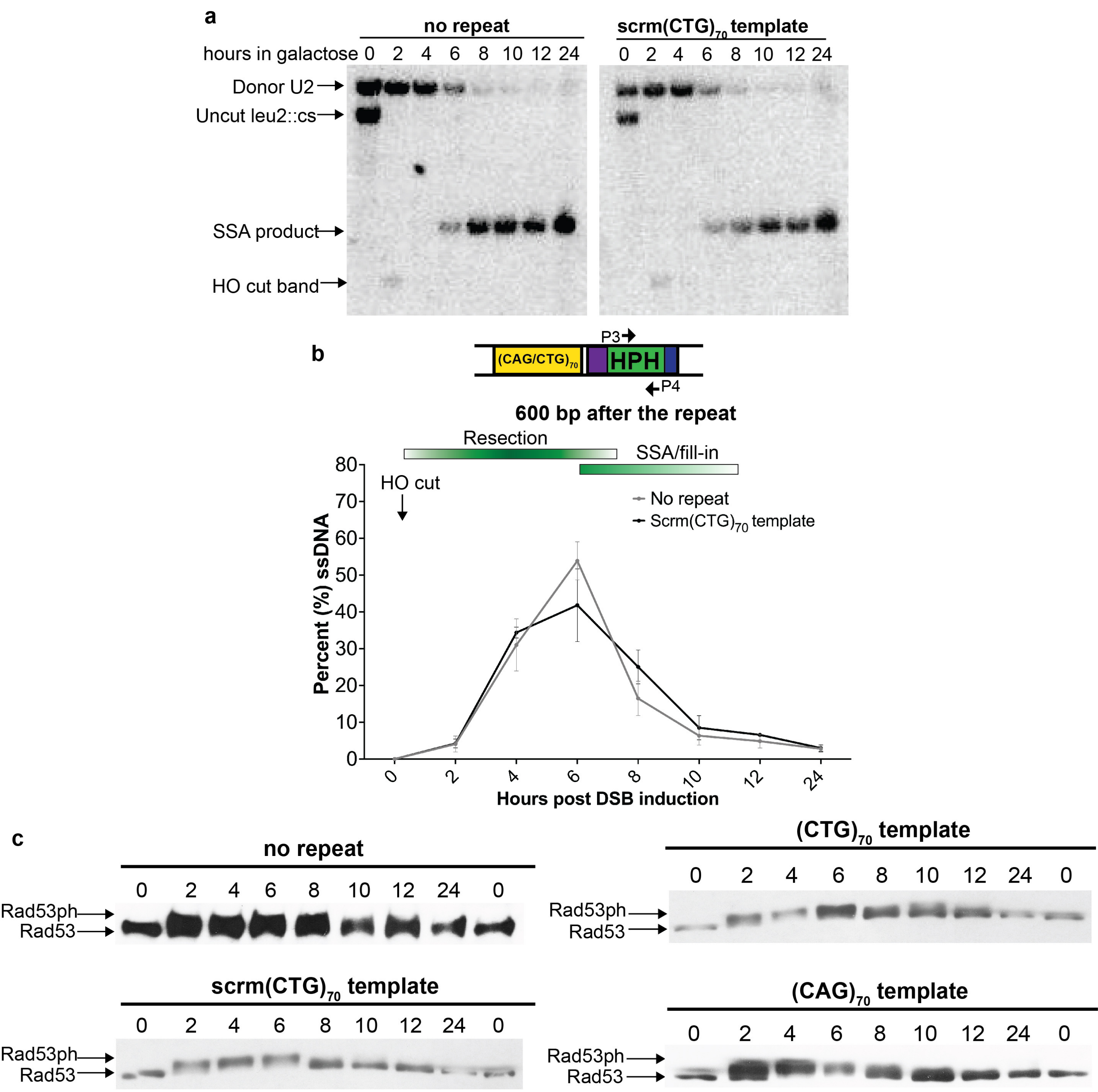

**Supplemental Figure 1: The scrm(CTG)$_{70}$ strain has similar repair kinetics to the no repeat control. a)** Southern blot analysis of no repeat (n=3) and scrm(CTG)70 (n=3) after addition of 2% galactose to induce a DSB within *LEU2*. Images are from the same Southern, scrm(CTG)70 is a different blot from the one in Figure 2a. **b)** Formation and disappearance of ssDNA after the repeat locus post DSB induction, using primers P3 and P4 in the no repeat (n=4) and scrm(CTG)$_{70}$ (n=3) strains. **c)** Protein lysates of no repeat (n=2), scrm(CTG)$_{70}$ (n=2), (CTG)$_{70}$ (n=2) and (CAG)$_{70}$ (n=2) repeat strains were analyzed by Western blot for hyper-phosphorylated Rad53 at indicated timepoints post DSB induction.

### Supplemental Figure 2

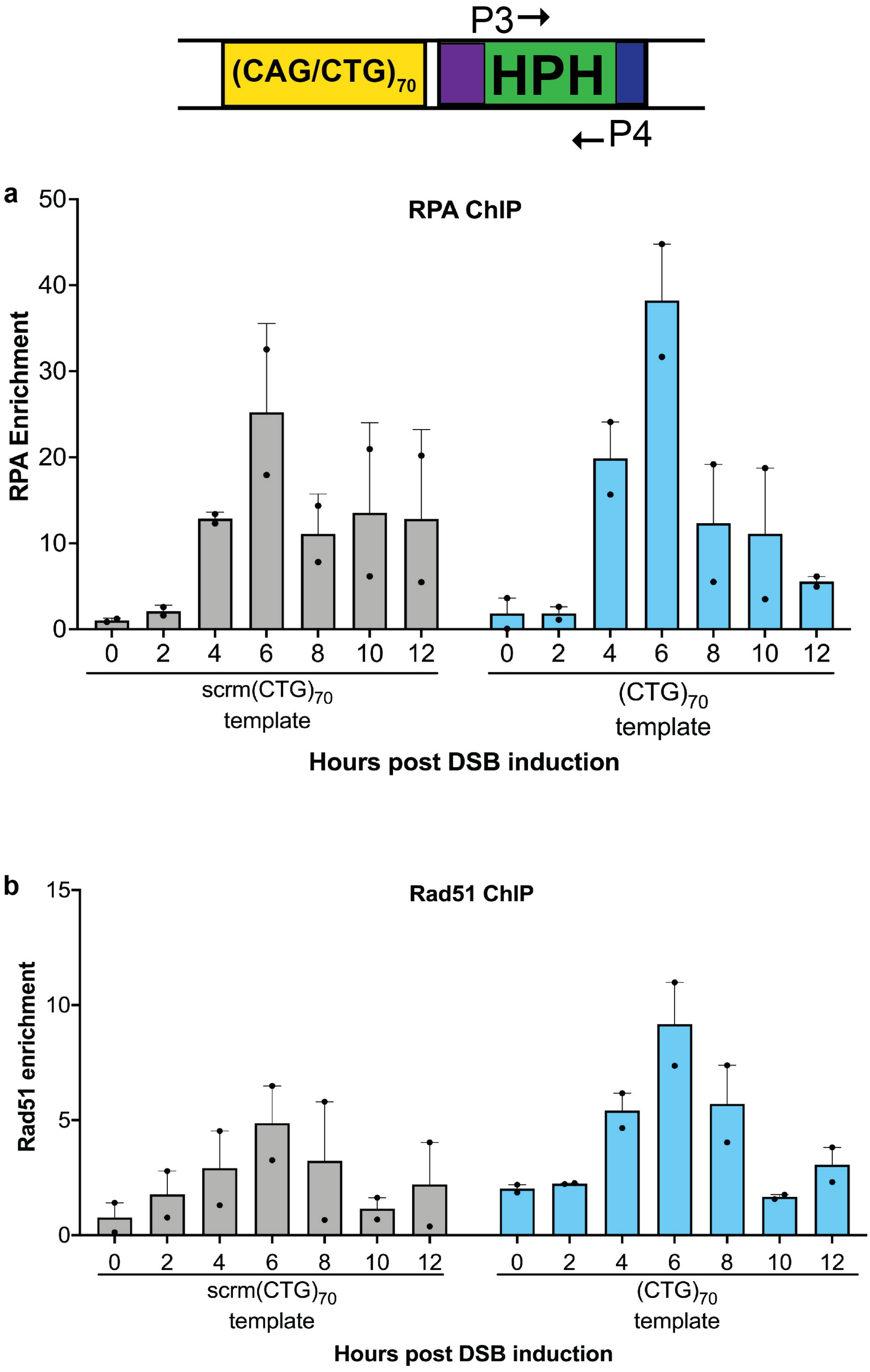

**Supplemental Figure 2: Enrichment of RPA and Rad51 on scrm$(CTG)_{70}$ and $(CTG)_{70}$ templates. a)** Enrichment of RPA adjacent to the repeat (P3-P4 amplicon) in scrm$(CTG)_{70}$ (n=2) and $(CTG)_{70}$ (n=2) template strains following DSB induction. Graph shows mean ± SEM. **b)** Enrichment of Rad51 adjacent to the repeat (P3-P4 amplicon) in scrm$(CTG)_{70}$ (n=2) and $(CTG)_{70}$ (n=2) template strains following DSB induction. Graph shows mean ± SEM.

### Supplemental Figure 3

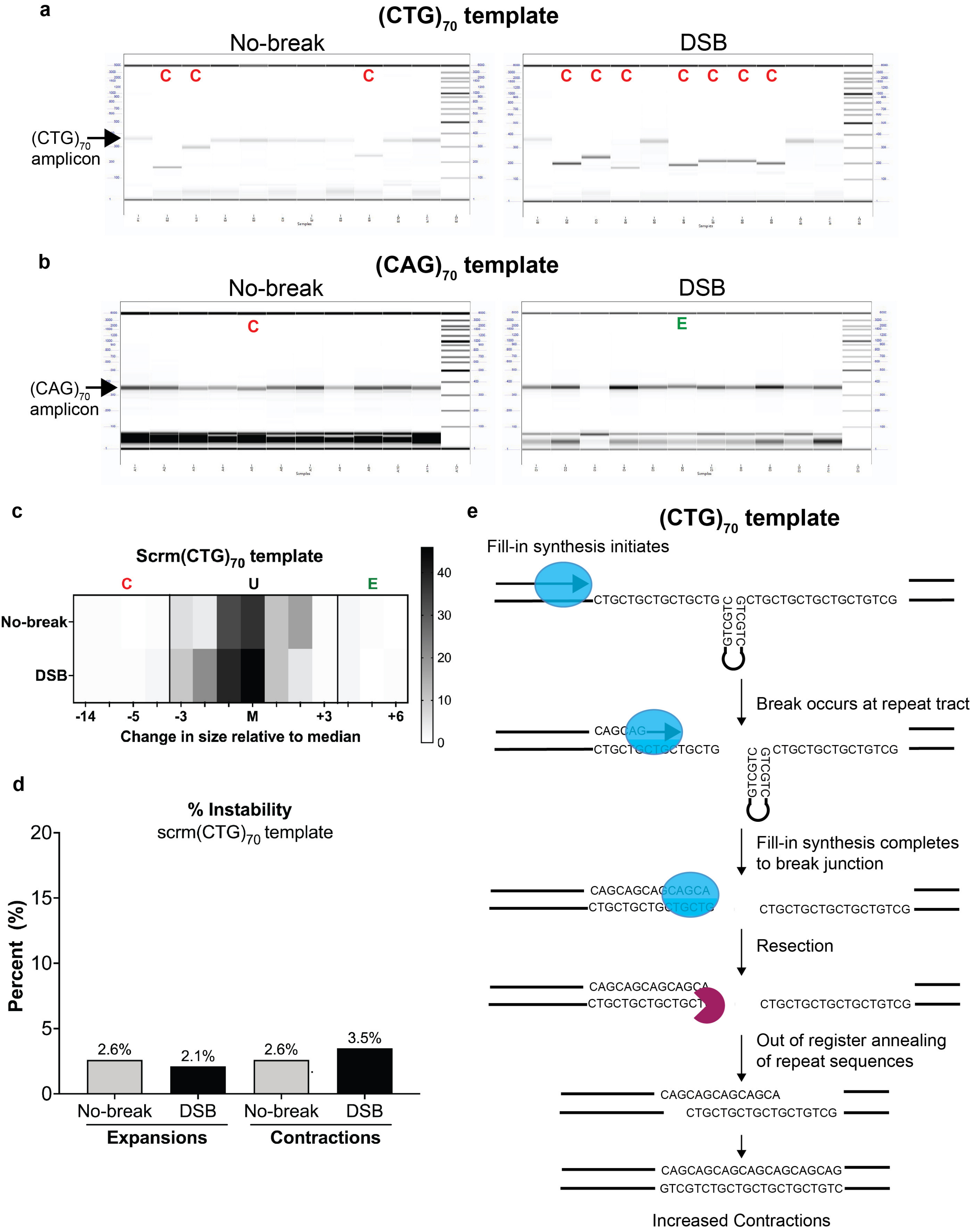

**Supplemental Figure 3: Visualization of $(CAG)_{70}$ and $(CTG)_{70}$ sizes and quantification of scrm$(CTG)_{70}$ instability. a)** Capillary gel electrophoresis images of $(CTG)_{70}$ repeat tract amplification in both the no-break and DSB conditions. Arrow points to unchanged size, C represents contracted samples. **b)** Capillary gel electrophoresis images of $(CAG)_{70}$ repeat tract amplification in both the no-break and DSB conditions . Arrow points to unchanged size, C represents contracted samples while E represents expanded samples. **c)** Heat map of amplicon sizes for the scrm$(CTG)_{70}$ sequence. Abbreviations are the same as Figure 3a. Total number of PCR reactions represented: no-break n=115, break n=141. **d)** Quantification of expansions and contractions of the scrm$(CTG)_{70}$ sequence. For the scrm$(CTG)_{70}$, no-break and DSB conditions showed no significant difference in expansions or contractions. **e)** Model for repeat contractions due to ssDNA breaks at the $(CTG)_{70}$ template. Resection and annealing of the U2 sequences occur unimpeded in the $(CTG)_{70}$ template strains. The ssDNA is left unprotected such that CTG hairpins can form and are targets of nucleolytic cleavage. Polymerase fill-in occurs independent of the break. Once the filled in strand is double stranded, resection and out of register alignment at the repeat tract occurs resulting in contractions.

### Supplemental Figure 4

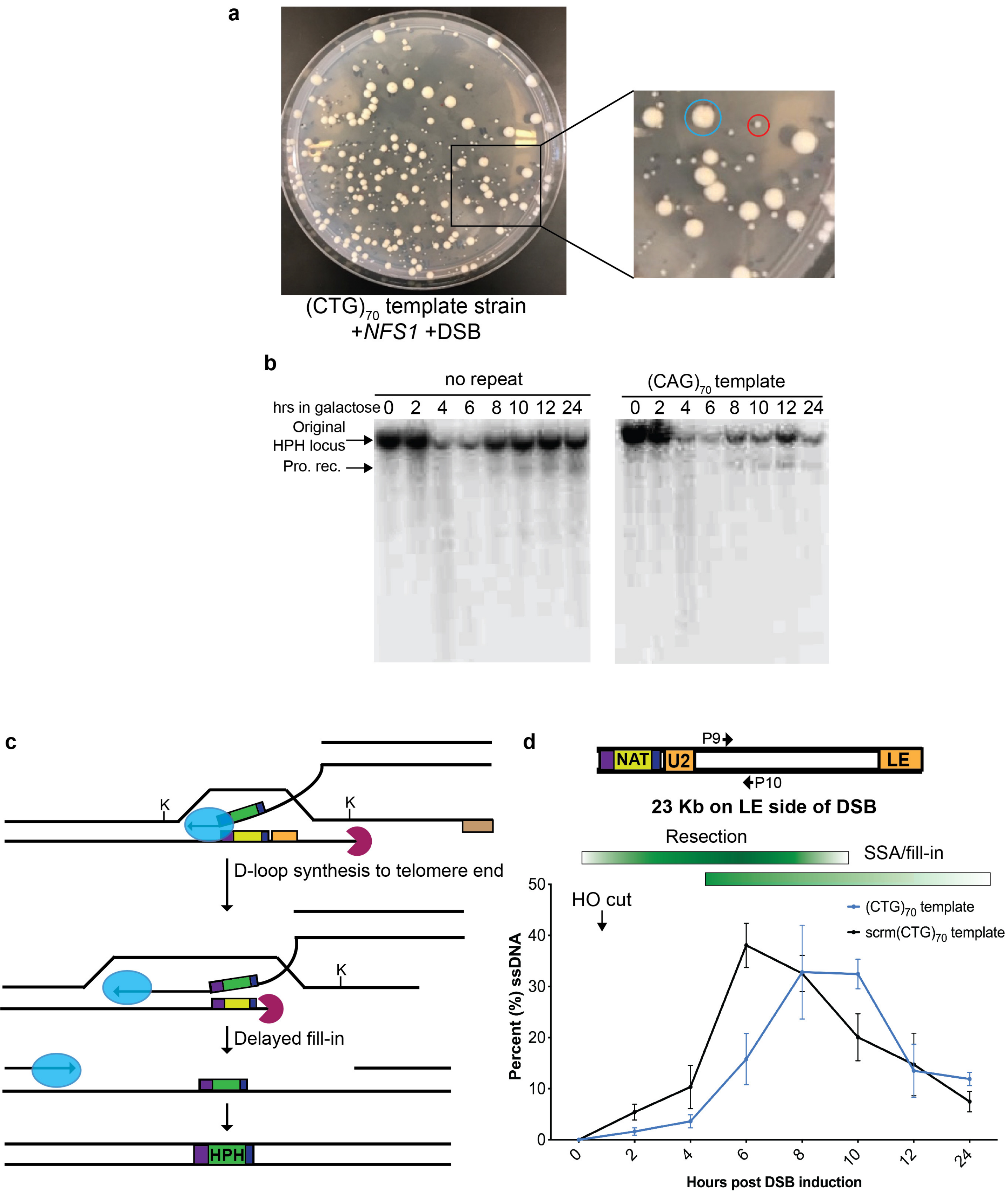

**Supplemental Figure 4: *NFS1* complemented colonies and resection on the other side of the DSB in the $(CTG)_{70}$ template strain. a)** There are two populations of resultant colonies from the $(CTG)_{70}$ strain containing the *NFS1* plasmid plated on break induction media. Colony circled in blue is a large colony, colony circled in red is a small colony. **b)** No tract and $(CAG)_{70}$ template Southern blots of KpnI digested DNA probed with a fragment to the *HPH* locus. Representative Southern shown; number of replicates: No repeat (n=2) and $(CAG)_{70}$ (n=2). **c)** Model for D-loop formation and filling in of BIR repair events in colonies with a $(CTG)_{70}$ ssDNA template that has broken (see Fig. 4a). BIR initiates from TEF promoter homology and D-loop synthesis occurs to the telomere end. Exonucleolytic degradation of the template is delayed as D-loop formation at the site of the TEF promoter occurs (see P9-10 amplicon data, Figure S4d). After leading strand BIR synthesis, filling in of the opposite strand is delayed in time (see P7-P8 amplicon data, Fig 4e). **d)** Resection and fill-in kinetics of strains with a $(CTG)_{70}$ repeat template is impaired on the other (U2) side of the DSB. Percent ssDNA from DSB induced strains 15 kb to the left of the DSB was determined as in Figure 2b. Number of replicates: scrm$(CTG)_{70}$ (n=3), $(CTG)_{70}$ (n=3).

Supplemental Figure 5

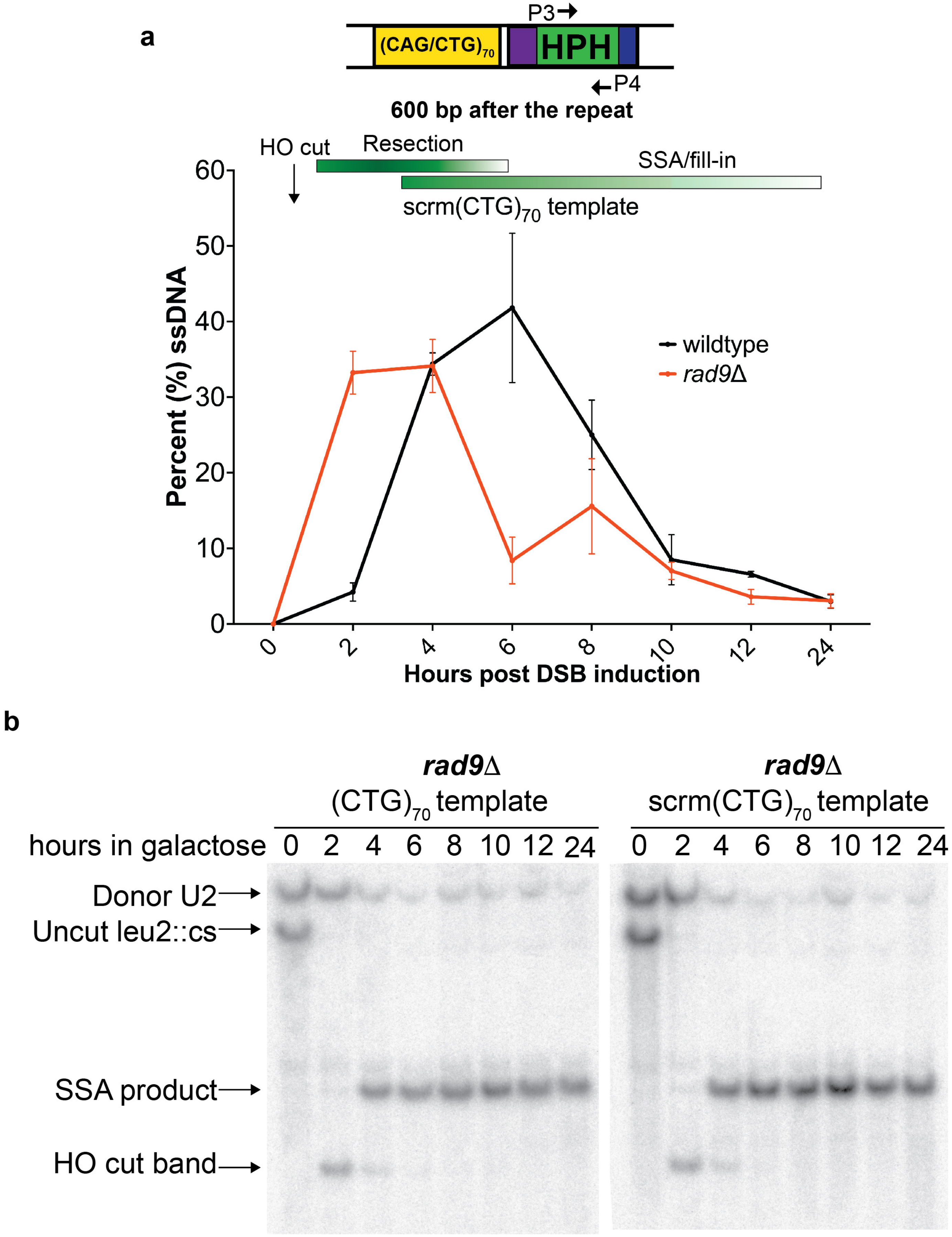

**Supplemental Figure 5: Deletion of Rad9 speeds up resection and repair. a)** Resection and fill-in kinetics of the scrm$(CTG)_{70}$ template construct in wildtype and *rad9Δ* strains. Percent ssDNA after the repeat locus from DSB induced strains was determined as in Figure 2b. Number of replicates: scrm$(CTG)_{70}$ (n=3), *rad9Δ* scrm$(CTG)_{70}$ (n=3). **b)** Southern blot analysis after addition of 2% galactose to induce a DSB within *LEU2* of the scrm$(CTG)_{70}$ and $(CTG)_{70}$ template constructs in *rad9Δ* mutants. Representative Southern shown; number of replicates: *rad9Δ* scrm$(CTG)_{70}$ (n=3), *rad9Δ* $(CTG)_{70}$ (n=3) and *rad9Δ* $(CAG)_{70}$ (n=2).

Supplemental Figure 6

a

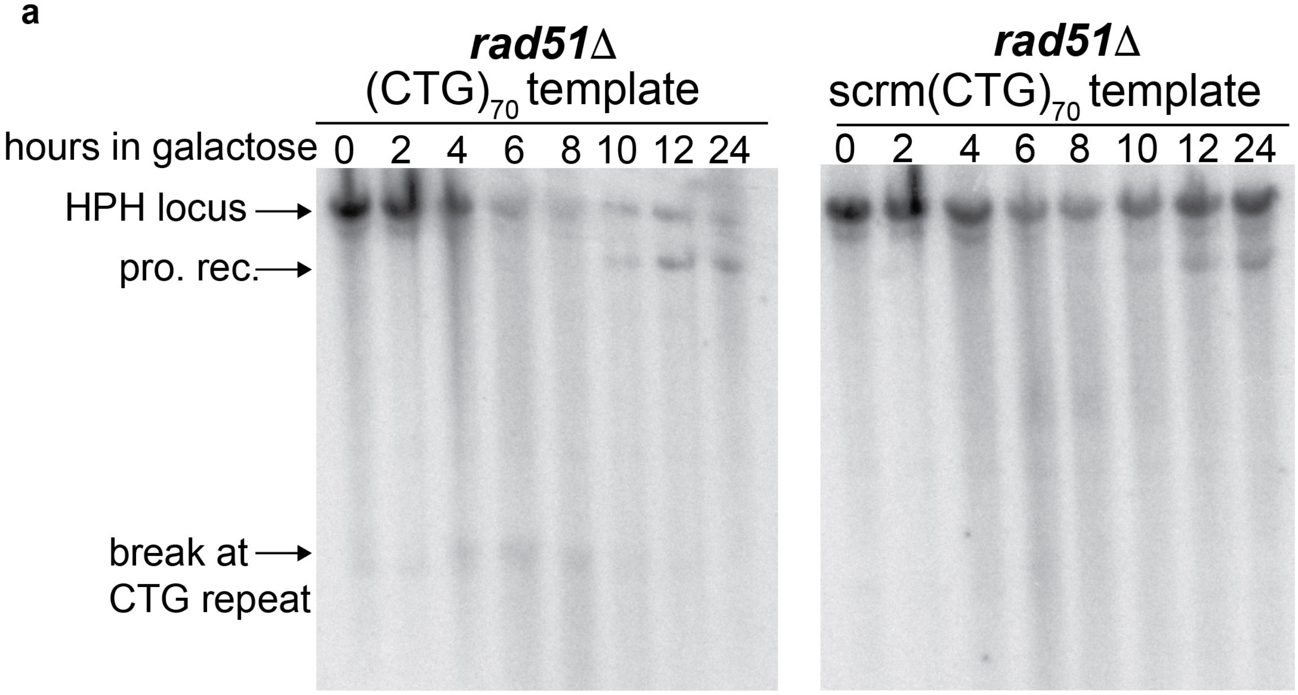

b

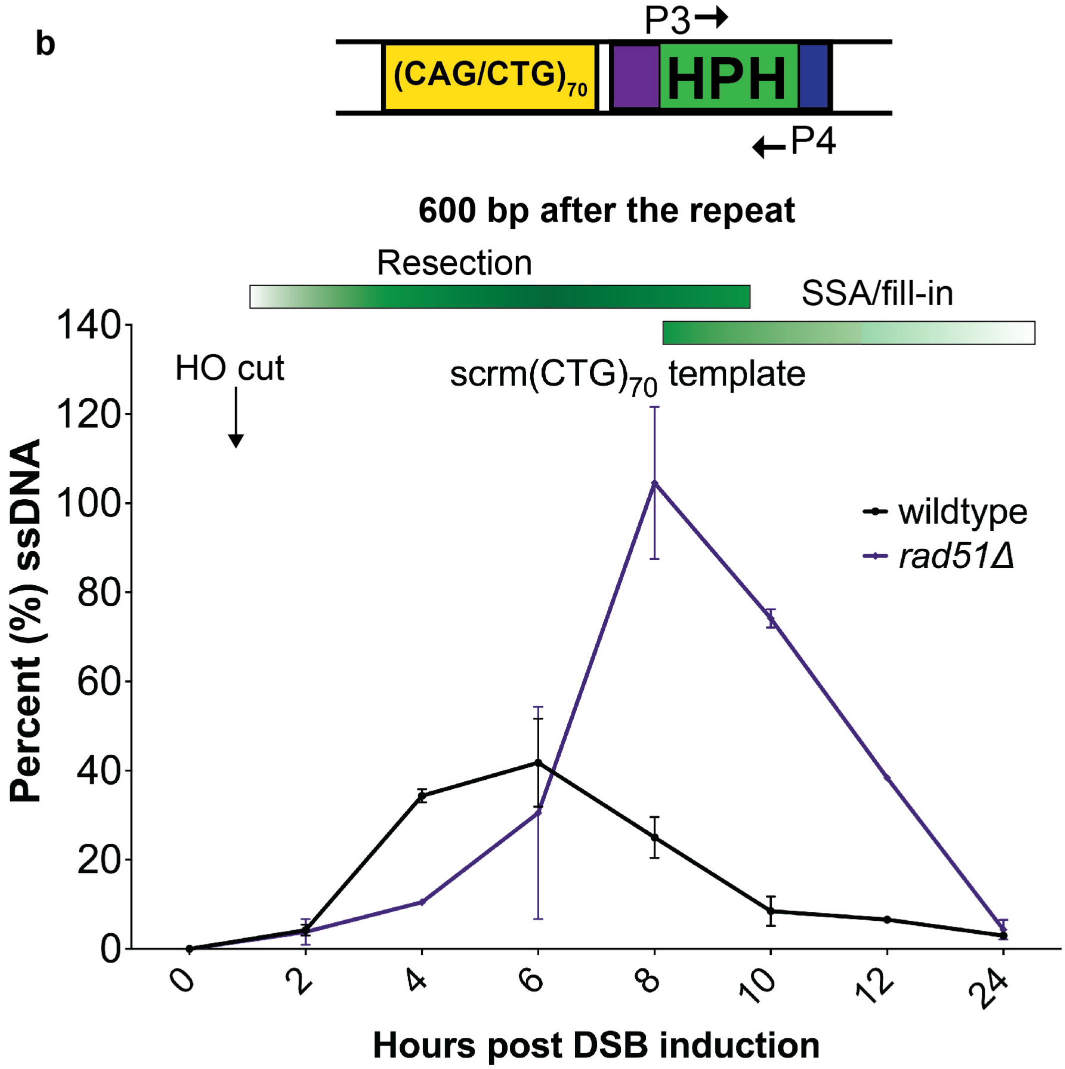

**Supplemental Figure 6: Deletion of Rad51 delays BIR repair and resection through the repeat tract. a)** Kinetic Southern blots (Figure 6e) of *rad51*Δ mutants in the scrm$(CTG)_{70}$ (n=2) and $(CTG)_{70}$ (n=2) strains were probed with a fragment to the *HPH* locus 600 bp downstream of the repeat tract. Representative Southern shown. **b)** Resection and fill-in kinetics of the scrm$(CTG)_{70}$ template construct in wildtype (n=3) and *rad51*Δ (n=2) strains. Percent ssDNA 600 bp after the repeat locus after DSB induction was determined as in Figure 3b.

**Supplemental Methods:**

**Rad53 Phosphorylation-** Time course collection was done at the same time as cells were collected for the kinetic Southern blot. For each timepoint, 9.25 x $10^7$ cells were collected. Protein was extracted via TCA method as previously described[1]. Westerns were probed with anti-Rad53 antibody (1:1000; mouse monoclonal EL7.E1 Abcam Cat#AB166859).

1 Foiani, M., Marini, F., Gamba, D., Lucchini, G. & Plevani, P. The B subunit of the DNA polymerase alpha-primase complex in Saccharomyces cerevisiae executes an essential function at the initial stage of DNA replication. *Mol Cell Biol* **14**, 923-933, doi:10.1128/mcb.14.2.923-933.1994 (1994).

| Table 1: Strains, Plasmids, Primers used | | |
|---|---|---|
| **Parent strain** | | |
| YMV80 | MATΔ::hisG hmlΔ::ADE1 hmrΔ::ADE1 ade1 lys5 ura3-52 trp1Δ HO ade3::GAL-HO leu2::cs | Vaze et al, 1999 |
| no repeat | same as YMV80, but ilv6Δ::HPH | this study |
| **scrm(CTG)70 template strains** | | |
| CFY4765, 4766 | same as YMV80, but ilv6Δ::(scrmCTG)70-HPH | this study |
| CFY5529, 5530 | same as parent 4797; rad9::trp1 | this study |
| CFY4946, 4947 | same as parent 4797; rad51::kanMX | this study |
| **(CAG)70 template strains** | | |
| CFY4017,4018,4019,4020 | same as YMV80, but ilv6Δ::(CAG)70-HPH | this study |
| **(CTG)70 template strains** | | |
| CFY4797, 4798 | same as YMV80, but ilv6Δ::(CTG)70-HPH | this study |
| CFY5351, 5352 | same as parent 4797; rad9::trp1 | this study |
| CFY4948, 4949, 4950 | same as parent 4797; rad51::kanMX | this study |
| **Plasmids** | | |
| pCF390 | (CTG)70-HPH | Su et al, 2015 |
| pCF590 | (CAG)70-HPH | this study |
| pCF722, 723 | scrm(CTG)70-HPH | this study |
| pCF187 | pRS414 | |
| pCF582, 583 | pRS414+NFS1 | this study |
| **Primers** | | |
| P1 - for scrm (CTG)70 and (CTG)70 templates forward | 5' CCC AGG CCT CCA GTT TGC 3' | |
| P2 - for scrm (CTG)70 and (CTG)70 templates reverse | 5’ TAA TAC GAC TCA CTA TAG GG 3’ | |
| P1 - for (CAG)70 templates forward | 5' CCG CCA GCT GAA GCT TGA AT 3' | |
| P1 - for (CAG)70 templates reverse | 5' CAG TTT GCC CAT CCA CGT CA 3' | |
| P3 - HPH locus forward | 5' AAA TAG CTG CGC CGA TGG TTT C 3' | |
| P3 - HPH locus forward (ChIP only) | 5' GAT TCC GGA AGT GCT TGA C 3' | |
| P4 - HPH locus reverse | 5' CAG CGA TCG CAT CCA TGG CCT C 3' | |
| P5 - BUD3 locus forward | 5’ ACT GCT GAA TTT CCG GTG GA 3’ | |
| P6 - BUD3 locus reverse | 5’ TCT GAG CAT TTC CTA ACA CGG T 3’ | |
| P7 - LDB16 locus forward | 5' CCA AAA CTC ATT CGT CAC CA 3' | |
| P8 - LDB16 locus reverse | 5' ACA TGG GTT ACT GGC AGA AA 3' | |
| P9 - RNQ1 locus forward | 5' GCT TTG GCG TCT TTG GCT TC 3' | |
| P10 - RNQ1 locus reverse | 5' CTC CAA AGG AGG AAC CAC CG 3' | |
| ACT1 forward | 5' TCC AGA TGG TCA AGT CAT CA 3' | |
| ACT1 reverse | 5' TCG GCA ATA CCT GGG AAC AT 3' | |

Table 2: Percent viability

| | Parent | No repeat | scrm(CTG)70 template | | | (CAG)70 template | (CTG)70 template | | | | |
|---|---|---|---|---|---|---|---|---|---|---|---|
| | | | wildtype | *rad9*Δ | *rad51*Δ | wildtype | wildtype | +TRP1 vector | +NFS1 | *rad9*Δ | *rad51*Δ |
| | 93.2 | 88.2 | 71.5 | 84.2 | 71.6 | 62.3 | 20.9 | 20.7 | 68.4 | 70.7 | 9.2 |
| | 69.6 | 78.2 | 73.7 | 77.1 | 60.5 | 51.4 | 11.7 | 37.4 | 50 | 77.9 | 7.7 |
| | 92.6 | 81.5 | 82.7 | 70.4 | 60.9 | 85.6 | 30.1 | 29.5 | 82.5 | 87 | 8.9 |
| | 76.4 | 73.8 | 82.6 | 77.4 | 54.8 | 76 | 17.8 | 35 | 45 | 58 | 10.9 |
| | 72.2 | 94 | 80.7 | 80 | 66.9 | 66.7 | 19.3 | 35.7 | 60.2 | 50 | 9.1 |
| | | 69.2 | 86.1 | 79 | 60.9 | 77.3 | | 25.3 | 54.5 | | 9.6 |
| | | 77.6 | 87.5 | | 54.8 | 79.6 | | 23.4 | 58.8 | | 10.6 |
| | | 73.3 | 78.5 | | 66.9 | 80.1 | | 19.4 | 55.4 | | 5.5 |
| | | 82.9 | 82.7 | | | 87 | | 33.7 | 57.4 | | 10.5 |
| | | 81.6 | 71 | | | 83.9 | | | 90 | | 18.5 |
| | | | 70.6 | | | 83.7 | | | | | 9.9 |
| | | | 88 | | | 79 | | | | | 10.4 |
| | | | | | | 89.8 | | | | | |
| | | | | | | 84.4 | | | | | |
| | | | | | | 71.3 | | | | | |
| | | | | | | 62.6 | | | | | |
| | | | | | | 75.7 | | | | | |
| | | | | | | 56.8 | | | | | |
| | | | | | | 74 | | | | | |
| **Average** | 80.8 | 80.0 | 79.6 | 78.0 | 62.2 | 75.1 | 20.0 | 28.9 | 62.2 | 68.7 | 10.1 |
| **# of replicates** | 5 | 10 | 12 | 6 | 8 | 19 | 5 | 9 | 10 | 5 | 12 |
| **Std. Dev** | 11.3 | 7.3 | 6.5 | 4.5 | 6.0 | 10.7 | 6.6 | 6.9 | 14.2 | 14.9 | 3.0 |
| **SEM** | 5.1 | 2.3 | 1.9 | 1.8 | 2.1 | 2.5 | 2.2 | 2.3 | 4.5 | 6.7 | 0.9 |
| **P-value to parent*** | | 0.87 | 0.79 | | | 0.31 | <0.0001 | | | | |
| **P-value to scrm(CTG 70)*** | | | | 0.59 | 0.00 | 0.20 | <0.0001 | | | | |
| **P-value to (CTG)70*** | | | | | | | | 0.01 | <0.0001 | <0.0001 | 0.0002 |
| **P-value to (CTG)70+ vector*** | | | | | | | | | <0.0001 | | |

* Students t-test

**Table 3: Listed experimental values for resection assays for all loci**

**HPH locus (P3 & P4)**

| Hour | No tract Ex. 1 | No tract Ex. 2 | No tract Ex. 3 | No tract Ex.4 | No tract Average | scrm(CTG)70 Ex. 1 | scrm(CTG)70 Ex. 2 | scrm(CTG)70 Ex. 3 | scrm(CTG)70 Average |
|---|---|---|---|---|---|---|---|---|---|
| 0 | 0 | 0 | 0 | 0 | **0** | 0 | 0 | 0 | **0** |
| 2 | 1.9 | 1.0 | 10.4 | 3 | **4.1** | 4.5 | 6.2 | 2.0 | **4.2** |
| 4 | 20.1 | 21.6 | 50.8 | 31.6 | **31.0** | 35.6 | 31.4 | 36.1 | **34.4** |
| 6 | 60.6 | 40.1 | 63.0 | 51.8 | **53.9** | 22.2 | 49.5 | 53.7 | **41.8** |
| 8 | 8.5 | 17.1 | 29.4 | 11 | **16.5** | 32.0 | 16.3 | 26.7 | **25.0** |
| 10 | 1.8 | 6.5 | 13.5 | 3.6 | **6.3** | 1.9 | 12.0 | 11.6 | **8.5** |
| 12 | 7.0 | 2.4 | 9.0 | 1 | **4.8** | 6.4 | 7.3 | 6.0 | **6.6** |
| 24 | 5.3 | 1.4 | 3.1 | 1.4 | **2.8** | 1.5 | 4.6 | 2.8 | **3.0** |

| Hour | $(CTG)_{70}$ template Ex. 1 | Ex. 2 | Ex. 3 | Ex.4 | Average | $(CAG)_{70}$ template Ex. 1 | Ex. 2 | Ex. 3 | Average |
|---|---|---|---|---|---|---|---|---|---|
| 0 | 0 | 0 | 0 | 0 | **0** | 0 | 0 | 0 | **0** |
| 2 | 15.9 | 6.7 | 2.2 | 1.8 | **6.7** | 3.9 | 0.62 | 3.1 | **2.5** |
| 4 | 32.0 | 24.1 | 33.1 | 28.2 | **29.3** | 23.28 | 7.74 | 41.8 | **24.3** |
| 6 | 64.3 | 72.3 | 65.1 | 67.3 | **67.3** | 29.78 | 8.54 | 29.7 | **22.7** |
| 8 | 43.0 | 47.6 | 41.4 | 39.2 | **42.8** | 8.89 | 4.01 | 7.7 | **6.9** |
| 10 | 29.7 | 26.5 | 21.4 | 24.0 | **25.4** | 4.46 | 1.1 | 3.4 | **3.0** |
| 12 | 17.5 | 13.0 | 12.0 | 10.1 | **13.1** | 9.33 | 0.34 | 2.7 | **4.1** |
| 24 | 27.2 | 8.9 | 9.6 | 8.0 | **13.4** | 3.2 | 0.5 | 1.8 | **1.8** |

**BUD3 locus (P5 & P6)**

| Hour | No tract Ex. 1 | Ex. 2 | Ex. 3 | Average | Scrm $(CTG)_{70}$ orientation Ex. 1 | Ex. 2 | Ex. 3 | Average |
|---|---|---|---|---|---|---|---|---|
| 0 | 0 | 0 | 0 | **0** | 0 | 0 | 0 | **0** |
| 2 | 6.7 | 16.6 | 14 | **12.4** | 13.1 | 7.72 | 20.4 | **13.7** |
| 4 | 39.3 | 54.8 | 22.2 | **38.8** | 56.86 | 56.26 | 65.9 | **59.7** |
| 6 | 35.0 | 43.4 | 65.1 | **47.8** | 48.54 | 50.21 | 63.6 | **54.1** |
| 8 | 10.6 | 16.7 | 8.9 | **12.1** | 10.58 | 19.35 | 11.8 | **13.9** |
| 10 | 4.1 | 5.0 | 3.1 | **4.1** | 8.19 | 8.54 | 6.7 | **7.8** |
| 12 | 1.9 | 4.4 | 1.1 | **2.5** | 4.52 | 4.78 | 4.2 | **4.5** |
| 24 | 1.0 | 1.9 | 1.3 | **1.4** | 2.3 | 1.97 | 3.7 | **2.7** |

| Hour | $(CTG)_{70}$ Orientation Ex. 1 | Ex. 2 | Ex. 3 | Average | $(CAG)_{70}$ Orientation Ex. 1 | Ex. 2 | Average |
|---|---|---|---|---|---|---|---|
| 0 | 0.0 | 0.0 | 0.0 | **0** | 0 | 0 | **0** |
| 2 | 24.2 | 11.2 | 8.9 | **14.7** | 3.13 | 17.3 | **10.2** |
| 4 | 49.3 | 32.4 | 48.1 | **43.3** | 18.81 | 69.2 | **44.0** |
| 6 | 51.6 | 54.6 | 50.8 | **52.3** | 5.59 | 35.9 | **20.7** |
| 8 | 21.4 | 22.3 | 18.7 | **20.8** | 2.29 | 5.6 | **3.9** |
| 10 | 14.9 | 11.7 | 8.5 | **11.7** | 0.7 | 2.2 | **1.5** |
| 12 | 9.2 | 6.4 | 4.9 | **6.8** | 0.28 | 1.8 | **1.0** |
| 24 | 19.8 | 4.5 | 3.6 | **9.3** | 0.42 | 2 | **1.2** |

**YCL locus (P7 & P8)**

| Hour | Scrm $(CTG)_{70}$ orientation Ex. 1 | Ex. 2 | Average | $(CTG)_{70}$ template Ex. 1 | Ex. 2 | Ex. 3 | Ex.4 | Average |
|---|---|---|---|---|---|---|---|---|
| 0 | 0 | 0 | **0.0** | 0 | 0 | 0 | 0 | **0** |
| 2 | 3.67 | 1.48 | **2.6** | 1.7 | 3.8 | 1.3 | 1.0 | **2.0** |
| 4 | 9.53 | 8.75 | **9.1** | 7.8 | 4.7 | 5.7 | 6.3 | **6.1** |
| 6 | 18.89 | 16.39 | **17.6** | 26.5 | 28.0 | 27.0 | 20.7 | **25.5** |
| 8 | 6.89 | 12.44 | **9.7** | 17.2 | 21.9 | 18.5 | 19.1 | **19.1** |
| 10 | 6.85 | 5.86 | **6.4** | 14.7 | 11.5 | 11.3 | 8.3 | **11.5** |
| 12 | 3.34 | 2.18 | **2.8** | 7.8 | 6.3 | 5.1 | 5.3 | **6.1** |
| 24 | 1.87 | 0.93 | **1.4** | 5.0 | 2.1 | 3.8 | 2.9 | **3.4** |

**RNQ locus (P9 & P10)**

| Hour | Scrm $(CTG)_{70}$ orientation Ex. 1 | Ex. 2 | Ex. 3 | Average | $(CTG)_{70}$ Orientation Ex. 1 | Ex. 2 | Ex. 3 | Ex.4 | Average |
|---|---|---|---|---|---|---|---|---|---|
| 0 | 0 | 0 | 0 | **0.0** | 0 | 0 | 0 | 0 | **0.0** |
| 2 | 7.1 | 6.8 | 2.3 | **5.4** | 2.3 | 0.5 | 3.3 | 0.3 | **1.6** |
| 4 | 13.9 | 15.2 | 1.9 | **10.3** | 3.8 | 4.2 | 6.3 | 0.1 | **3.6** |
| 6 | 41.1 | 43.6 | 29.5 | **38.1** | 18.3 | 16.9 | 25.9 | 2.0 | **15.8** |
| 8 | 34 | 37.9 | 25.8 | **32.6** | 53.1 | 39.2 | 29.7 | 9.3 | **32.8** |
| 10 | 25.4 | 23.9 | 10.9 | **20.1** | | 31.8 | 37.8 | 27.8 | **32.4** |
| 12 | 21.3 | 20.4 | 2.5 | **14.7** | 23.5 | 4.9 | 21.5 | 4.2 | **13.5** |
| 24 | 9 | 9.9 | 3.5 | **7.5** | 9.9 | | 11.5 | 14.3 | **11.9** |

**Mutant data: HPH locus (P3 & P4)**

| Hour | *rad9Δ* scrm$(CTG)_{70}$ template Ex. 1 | Ex. 2 | Ex. 3 | Average | *rad51Δ* scrm(CTG)70 template Ex. 1 | Ex. 2 | Average |
|---|---|---|---|---|---|---|---|
| 0 | 0 | 0 | 0 | **0.0** | 0 | 0 | **0** |
| 2 | 35.4 | 27.6 | 36.7 | **33.2** | 0.9 | 6.7 | **3.8** |
| 4 | 34 | 28.1 | 40.3 | **34.1** | 10.0 | 11.0 | **10.5** |
| 6 | 14.5 | 4.4 | 6.3 | **8.4** | 54.4 | 6.7 | **30.5** |
| 8 | 24.7 | 3.5 | 18.5 | **15.6** | 121.6 | 87.5 | **104.6** |
| 10 | 5.9 | 5.9 | 9.3 | **7.0** | 72.1 | 76.2 | **74.1** |
| 12 | 3.1 | 2.2 | 5.5 | **3.6** | 37.9 | 38.9 | **38.4** |
| 24 | 1.8 | 2.5 | 4.9 | **3.1** | 2.2 | 6.5 | **4.4** |

| Hour | *rad9Δ* $(CTG)_{70}$ template Ex. 1 | Ex. 2 | Ex. 3 | Average | *rad51Δ* (CTG)70 template Ex. 1 | Ex. 2 | Average |
|---|---|---|---|---|---|---|---|
| 0 | 0 | 0 | 0 | **0.0** | 0 | 0 | **0.0** |
| 2 | 63.3 | 36.0 | 50.8 | **50.0** | 2.01 | 3.7 | **2.9** |
| 4 | 40.1 | 30.1 | 54.9 | **41.7** | 7.28 | 3 | **5.1** |
| 6 | 7.8 | 7.0 | 13.3 | **9.4** | 57.71 | 27.8 | **42.8** |
| 8 | 4.5 | 4.9 | 6.1 | **5.2** | 88.57 | 83.3 | **85.9** |
| 10 | 5.1 | 2.9 | 5.2 | **4.4** | 50.33 | 94.3 | **72.3** |
| 12 | 3.8 | 2.7 | 4.8 | **3.8** | 66.22 | 107.4 | **86.8** |
| 24 | 2.8 | 2.7 | 2.3 | **2.6** | 3.15 | 6.2 | **4.7** |

**Table 4: Instability frequency**

| (CAG)70 template strain | | | | | | | | | | | | | |
|---|---|---|---|---|---|---|---|---|---|---|---|---|---|
| | **Expansions** | | | | | | | **Contractions** | | | | | |
| | no-break expansions (total) | Percent | | DSB expansions (total) | Percent | pvalue DSB to no-break | | no-break contractions (total) | Percent | | DSB contractions (total) | Percent | pvalue DSB to no-break |
| Wildtype (CAG)70 | 9 (261) | 3.4% | | 51 (273) | 18.70% | 0.0001 | | 50 (261) | 19.2% | | 15 (273) | 5.5% | 0.0001 |

| (CTG)70 template strain | | | | | | | | | | | | | | |
|---|---|---|---|---|---|---|---|---|---|---|---|---|---|---|
| | **Expansions** | | | | | | | **Contractions** | | | | | | |
| | no-break expansions (total) | Percent | pvalue to WT no break | DSB expansions (total) | Percent | pvalue DSB to no-break | pvalue to WT DSB | no-break contractions (total) | Percent | pvalue to WT no break | DSB contractions (total) | Percent | pvalue DSB to no-break | pvalue to WT DSB |
| Wildtype (CTG)70 | 2 (120) | 2.5% | | 2 (119) | 1.7% | 1 | | 58 (120) | 48.3% | | 93(119) | 78.2% | 0.03 | |
| *rad9*Δ | 3 (163) | 1.8% | 1 | 2 (163) | 1.2% | 1 | 1 | 49 (163) | 27.4% | 0.04 | 65 (163) | 41.7% | 0.23 | 0.001 |
| *rad51*Δ | 2(119) | 1.7% | 1 | 0 (119) | 0 | 0.50 | 0.50 | 40(119) | 33.6% | 0.15 | 116(119) | 97.5% | 0.0001 | 0.26 |

| scrm(CTG)70 template strain | | | | | | | | | | | | | |
|---|---|---|---|---|---|---|---|---|---|---|---|---|---|
| | **Expansions** | | | | | | | **Contractions** | | | | | |
| | no-break expansions (total) | Percent | | DSB expansions (total) | Percent | pvalue DSB to no-break | | no-break contractions (total) | Percent | | DSB contractions (total) | Percent | pvalue DSB to no-break |
| Wildtype scrm(CTG)70 | 3(115) | 2.6% | | 3 (141) | 2.1% | 1 | | 3(115) | 2.6% | | 5(141) | 3.5% | 0.7349 |

p values determined using Fisher's exact test

**Table 5: Fragment analyzer sizes called**

**(CTG)70 template**

| | | size in bp | no-break | DSB |
|---|---|---|---|---|
| contractions | | 144 | 0 | 1 |
| | | 156 | 1 | 0 |
| | | 161 | 1 | 0 |
| | | 162 | 0 | 1 |
| | | 163 | 0 | 2 |
| | | 164 | 0 | 1 |
| | | 165 | 1 | 0 |
| | | 168 | 0 | 4 |
| | | 173 | 0 | 1 |
| | | 174 | 0 | 3 |
| | | 175 | 1 | 0 |
| | | 177 | 0 | 1 |
| | | 179 | 0 | 1 |
| | | 180 | 1 | 1 |
| | | 181 | 2 | 1 |
| | | 182 | 1 | 1 |
| | | 186 | 1 | 0 |
| | | 187 | 1 | 6 |
| | | 190 | 0 | 2 |
| | | 192 | 0 | 3 |
| | | 193 | 2 | 3 |
| | | 195 | 0 | 1 |
| | | 196 | 0 | 3 |
| | | 198 | 2 | 0 |
| | | 199 | 0 | 2 |
| | | 200 | 0 | 3 |
| | | 203 | 0 | 1 |
| | | 204 | 0 | 1 |
| | | 205 | 1 | 1 |
| | | 206 | 0 | 3 |
| | | 208 | 1 | 0 |
| | | 209 | 0 | 1 |
| | | 210 | 0 | 1 |
| | | 211 | 1 | 0 |
| | | 214 | 2 | 0 |
| | | 215 | 0 | 3 |
| | | 216 | 0 | 2 |
| | | 218 | 4 | 3 |
| | | 221 | 0 | 1 |
| | | 223 | 1 | 0 |
| | | 224 | 2 | 2 |
| | | 227 | 3 | 6 |
| | | 228 | 3 | 6 |
| | | 229 | 0 | 1 |
| | | 230 | 1 | 2 |
| | | 231 | 1 | 0 |
| | | 234 | 0 | 1 |
| | | 237 | 1 | 0 |
| | | 238 | 1 | 1 |
| | | 239 | 0 | 1 |
| | | 240 | 1 | 1 |
| | | 242 | 1 | 1 |
| | | 294 | 0 | 1 |
| | | 252 | 1 | 0 |
| | | 253 | 0 | 1 |
| | | 254 | 1 | 0 |
| | | 256 | 1 | 0 |
| | | 258 | 0 | 1 |
| | | 260 | 1 | 0 |
| | | 261 | 1 | 0 |
| | | 262 | 1 | 0 |
| | | 264 | 1 | 0 |
| | | 265 | 1 | 0 |
| | | 266 | 1 | 0 |
| | | 272 | 0 | 1 |
| | | 273 | 0 | 1 |
| | | 280 | 0 | 1 |
| | | 287 | 1 | 0 |
| | | 295 | 0 | 1 |
| | | 296 | 1 | 0 |
| | | 306 | 1 | 0 |
| | | 308 | 0 | 1 |
| | | 321 | 1 | 0 |
| | | 322 | 1 | 0 |
| | | 343 | 3 | 2 |
| | | 344 | 1 | 0 |
| | | 345 | 0 | 1 |
| unchanged | | 347 | 5 | 0 |
| | | 348 | 6 | 0 |
| | | 349 | 14 | 3 |
| | Median | **350** | **14** | **10** |
| | | 351 | 11 | 4 |
| | | 352 | 7 | 6 |
| | | 353 | 2 | 1 |
| expansions | | 354 | 1 | 0 |
| | | 356 | 1 | 0 |
| | | 357 | 1 | 2 |
| | | **Total** | 118 | 117 |
| | | **Contractions** | | |
| | | Total | 56 | 91 |
| | | % | 47.5% | 77.8% |
| | | **Expansions** | | |
| | | Total | 3 | 2 |
| | | % | 2.5% | 1.7% |

**(CAG)70 template**

| | | size in bp | no-break | DSB |
|---|---|---|---|---|
| contractions | | 208 | 0 | 1 |
| | | 216 | 0 | 1 |
| | | 303 | 0 | 1 |
| | | 311 | 0 | 1 |
| | | 312 | 1 | 0 |
| | | 313 | 0 | 1 |
| | | 316 | 0 | 1 |
| | | 340 | 1 | 0 |
| | | 342 | 0 | 2 |
| | | 343 | 1 | 0 |
| | | 344 | 0 | 1 |
| | | 345 | 1 | 0 |
| | | 346 | 3 | 0 |
| | | 347 | 8 | 0 |
| | | 348 | 5 | 0 |
| | | 350 | 6 | 0 |
| | | 351 | 11 | 2 |
| | | 352 | 13 | 4 |
| unchanged | | 353 | 13 | 19 |
| | | 354 | 14 | 10 |
| | | 355 | 35 | 27 |
| | Median | **356** | **50** | **24** |
| | | **357** | **49** | **50** |
| | | 358 | 22 | 33 |
| | | 359 | 14 | 23 |
| | | 360 | 5 | 21 |
| expansions | | 361 | 3 | 26 |
| | | 362 | 4 | 11 |
| | | 363 | 0 | 8 |
| | | 364 | 1 | 4 |
| | | 365 | 0 | 1 |
| | | 366 | 0 | 1 |
| | | 440 | 1 | 0 |
| | | **Total** | 261 | 273 |
| | | **Contractions** | | |
| | | total | 50.00 | 15.00 |
| | | % | 19.2% | 5.5% |
| | | **Expansions** | | |
| | | total | 9 | 51 |
| | | % | 3.4% | 18.7% |

**scrm(CTG)70 template**

| | | size in bp | no-break | DSB |
|---|---|---|---|---|
| contractions | | 336 | 1 | 1 |
| | | 342 | 1 | 1 |
| | | 345 | 0 | 1 |
| | | 346 | 1 | 2 |
| unchanged | | 347 | 6 | 10 |
| | | 348 | 4 | 21 |
| | | 349 | 33 | 39 |
| | Median | **350** | **37** | **46** |
| | | 351 | 11 | 11 |
| | | 352 | 17 | 5 |
| | | 353 | 1 | 1 |
| expansions | | 354 | 2 | 2 |
| | | 355 | 1 | 0 |
| | | 356 | 0 | 1 |
| | | **Total** | 115 | 141 |
| | | **Contractions** | | |
| | | total | 3 | 5 |
| | | % | 2.60% | 3.50% |
| | | **Expansions** | | |
| | | total | 3 | 3 |
| | | % | 2.60% | 2.10% |

**Table 6: Enrichment of RPA and Rad51 by Chromatin Immunoprecipitation.**

| | RPA Enrichment at HPH locus (P3 & P4) relative to ACT1 | | | | | | | | | | | |
|---|---|---|---|---|---|---|---|---|---|---|---|---|
| | scrm(CTG)70 template | | | (CTG)70 template | | | *rad9* Δ (CTG)70 template | | | *rad51* Δ (CTG)70 template | | |
| **Hour** | **Ex. 1** | **Ex. 2** | **Average** | **Ex. 1** | **Ex. 2** | **Average** | **Ex. 1** | **Ex. 2** | **Average** | **Ex. 1** | **Ex. 2** | **Average** |
| **0** | 1.2 | 0.9 | **1.0** | 3.6 | 0.1 | **1.8** | 1.0 | 0.7 | **0.9** | 1.4 | 1.6 | **1.5** |
| **2** | 2.6 | 1.6 | **2.1** | 2.6 | 1.1 | **1.9** | 24.6 | 16.1 | **20.3** | 2.4 | 3.1 | **2.8** |
| **4** | 12.3 | 13.4 | **12.9** | 24.1 | 15.7 | **19.9** | 15.0 | 14.6 | **14.8** | 3.1 | 3.9 | **3.5** |
| **6** | 32.5 | 17.9 | **25.2** | 31.7 | 44.8 | **38.2** | 4.3 | 0.5 | **2.4** | 8.7 | 48.9 | **28.8** |
| **8** | 7.8 | 14.4 | **11.1** | 19.2 | 5.5 | **12.3** | 3.6 | 2.4 | **3.0** | 23.6 | 47.2 | **35.4** |
| **10** | 21.0 | 6.2 | **13.6** | 18.7 | 3.5 | **11.1** | | | | 29.3 | 4.4 | **16.9** |
| **12** | 20.2 | 5.5 | **12.8** | 5.0 | 6.1 | **5.6** | | | | 8.4 | 9.1 | **8.7** |

| | Rad51 Enrichment at HPH locus (P3 & P4) relative to ACT1 | | | | | | | | |
|---|---|---|---|---|---|---|---|---|---|
| | scrm(CTG)70 template | | | (CTG)70 template | | | *rad9* Δ (CTG)70 template | | |
| **Hour** | **Ex. 1** | **Ex. 2** | **Average** | **Ex. 1** | **Ex. 2** | **Average** | **Ex. 1** | **Ex. 2** | **Average** |
| **0** | 1.4 | 0.1 | **0.8** | 1.85 | 2.19 | **2.0** | 0.05 | 3.05 | **1.6** |
| **2** | 2.8 | 0.8 | **1.8** | 2.26 | 2.22 | **2.2** | 5.59 | 3.15 | **4.4** |
| **4** | 4.5 | 1.3 | **2.9** | 6.17 | 4.66 | **5.4** | 1.15 | 7.58 | **4.4** |
| **6** | 6.5 | 3.3 | **4.9** | 10.99 | 7.36 | **9.2** | 1.5 | 2.11 | **1.8** |
| **8** | 5.8 | 0.7 | **3.2** | 7.38 | 4.03 | **5.7** | 1.83 | 4.55 | **3.2** |
| **10** | 1.6 | 0.7 | **1.2** | 1.76 | 1.57 | **1.7** | | | |
| **12** | 4.0 | 0.4 | **2.2** | 3.81 | 2.31 | **3.1** | | | |
